# Evolutionarily divergent *Mycobacterium tuberculosis* CTP synthase filaments are under selective pressure

**DOI:** 10.1101/2024.07.25.605180

**Authors:** Eric M. Lynch, Yao Lu, Jin Ho Park, Lin Shao, Justin Kollman, E. Hesper Rego

## Abstract

The final and rate-limiting enzyme in pyrimidine biosynthesis, CTP synthase (CTPS), is essential for the viability of *Mycobacterium tuberculosis* and other mycobacteria. Its product, CTP, is critical for RNA, DNA, lipid and cell wall synthesis, and is involved in chromosome segregation. In various organisms across the tree of life, CTPS assembles into higher-order filaments, leading us to hypothesize that *M. tuberculosis* CTPS (mtCTPS) also forms higher-order structures. Here, we show that mtCTPS does assemble into filaments but with an unusual architecture not seen in other organisms. Through a combination of structural, biochemical, and cellular techniques, we show that polymerization stabilizes the active conformation of the enzyme and resists product inhibition, potentially allowing for the highly localized production of CTP within the cell. Indeed, CTPS filaments localize near the CTP-dependent complex needed for chromosome segregation, and cells expressing mutant enzymes unable to polymerize are altered in their ability to robustly form this complex. Intriguingly, mutants that alter filament formation are under positive selection in clinical isolates of *M. tuberculosis*, pointing to a critical role needed to withstand pressures imposed by the host and/or antibiotics. Taken together, our data reveal an unexpected mechanism for the spatially organized production of a critical nucleotide in *M. tuberculosis*, which may represent a vulnerability of the pathogen that can be exploited with chemotherapy.

## INTRODUCTION

CTP is critical to cellular function. In addition to being a core component of RNA and DNA, CTP is essential in phospholipid and cell wall biosynthesis as well as the segregation of chromosomes and low-copy-number plasmids during bacterial replication^1,2^. The rate-limiting step of CTP biosynthesis, the conversion of UTP to CTP, is catalyzed by CTPS. CTPS is a homotetramer, with each monomer comprising two domains, an amidoligase domain and a glutaminase domain joined by an alpha-helical linker ^3^. Ammonia generated via glutamine hydrolysis in the glutaminase domain is transferred to the amidologiase domain, where it is ligated to UTP in an ATP-dependent process, yielding CTP. CTPS undergoes feedback inhibition by CTP, which binds at two sites overlapping the UTP and ATP substrate-binding sites^4–7^. Substrate- and product-binding control a conformational cycle between active and inhibited states: UTP or CTP binding induces extension or compression of the tetramer interface, respectively. Further, UTP binding promotes rotation of the glutaminase domain towards the amidoligase domain, thus opening a tunnel to facilitate ammonia transfer between the two active sites^5,8,9^. GTP enhances CTPS activity by binding adjacent to this tunnel, preventing ammonia leakage through an opening at the junction between the glutaminase and amidoligase domains^5,10^. Notably, CTPS is therefore sensitive to levels of all four major ribonucleotides, highlighting its important regulatory role in nucleotide metabolism.

Polymerization of tetramers into filaments adds an additional layer of regulation to CTPS. There is remarkable diversity in the structure and function of CTPS filaments amongst different species. In all studied cases, however, filaments function to stabilize, prevent, or couple the highly-conserved conformational changes observed in CTPS tetramers. *E. coli* CTPS assembles into inactive filaments upon binding CTP, with tetramers interlocked via contacts in the linker and glutaminase domains such that transition to the active, UTP-bound conformation is sterically inhibited^8,11,12^. CTPS filaments from eukaryotes assemble in a different manner, with tetramers stacking via a eukaryotic-specific helical insert in the glutaminase domain. Human CTPS1—the proliferative isoform of the enzyme—forms hyper-active filaments in the presence of substrates^8^. By contrast, human CTPS2—the housekeeping isoform of the enzyme—can switch between active substrate-bound and inhibited product-bound filament conformations^9^. This has the effect of enhancing cooperativity, as cooperative conformational changes are no longer restricted to the closed symmetry of a tetramer, but can instead propagate along unbounded filaments. *Drosophila* and yeast CTPS have similarly been shown to form filaments with both substrate- and product-bound conformations^5,7^. Yeast CTPS has a different filament assembly interface and consequently distinct filament architectures; assembly into filaments prevents yeast CTPS from adopting a fully active conformation, resulting in reduced enzyme activity^7^.

Although their cellular function remains largely unknown, CTPS filaments appear in response to nutrient stress, at specific developmental stages, and in cancer cells, overall suggesting a role in adapting to conditions with particular metabolic needs ^13–17^. Indeed, yeast CTPS filaments are necessary for normal growth in both log phase and upon recovery from starvation^7^. In *Drosophila* early stage egg chambers, inhibiting CTPS polymerization leads to reduced egg production^18^. CTPS filaments also regulate cell curvature in *Caulobacter crescentus,* in a manner independent of CTPS catalytic activity^19^.

CTPS inhibition is a target for the treatment of both infectious and noncommunicable human diseases. For example, a variety of small molecules have been shown to target *M. tuberculosis* CTPS. The *pyrG* gene encoding CTPS is essential in *M. tuberculosis* ^20,21^, and inhibitors targeting CTPS prevent growth of the the pathogen ^22,23^. Here, we show that, as in other organisms, *M. tuberculosis* CTPS forms filaments, though their structure differs dramatically from other known CTPS filaments, with subunits packing in a different orientation. Filaments form in defined subcellular locations during rapid exponential growth and disappear during growth inhibition. Biochemical assays reveal that mtCTPS filaments stabilize the active enzyme conformation and resist feedback inhibition by CTP. Further, mutations that affect filament formation and enzyme kinetics are positively selected for in *M. tuberculosis* clinical isolates, suggesting a functional role in mycobacterial physiology and pathogenesis. Finally, we show that small molecules can specifically inhibit mycobacterial CTPS but not the human isoforms, solidifying CTPS as a bona fide drug target for *M. tuberculosis*. Together, our data suggest that filamentation of this critical enzyme arose independently through evolution and adds to the known functions performed by enzymatic filaments throughout the tree of life.

## RESULTS

### Mycobacterial CTPS forms filaments during exponential growth

As CTPS forms filaments in diverse organisms, we asked if mycobacterial CTPS also existed in higher-order structures. To investigate this, we generated a strain of *Mycobacterium smegmatis* expressing CTPS-eGFP at the chromosomal locus, ensuring native expression levels and regulation. CTPS is highly conserved throughout the mycobacterial genus, and *M. smegmatis* and *M. tuberculosis* CTPS share 87% amino acid identity. *pyrG* (the gene encoding CTPS) is essential for the viability of *M. smegmatis* and *M. tuberculosis* ^20,21,24^. Thus, the normal growth rate of this strain indicates that CTPS-eGFP is fully functional (Fig. S1). Imaging by fluorescence live-cell microscopy revealed that CTPS-eGFP does indeed exist in discrete puncta. The puncta were mobile but were often found near the poles, with new puncta frequently emerging at septa before division (Fig. 1a). To gain a clearer understanding of the puncta’s shape and location in the cell, we performed two-color live-cell 3D structured-illumination (3D-SIM), counter-staining with a fluorescent d-amino acid that incorporates into the bacterial cell wall. Using this super-resolution technique, we observed that CTPS-eGFP was not isotropic in shape, but instead most puncta resolved into small linear structures (Fig. 1b,c). Visualizing CTPS-eGFP localization in the third dimension, along the cross-section of the bacterium, revealed that CTPS-eGFP was almost always confined to the periphery of the bacterium, presumably next to the plasma membrane (Fig. 1c). Performing two-color 3D-SIM at fast timescales showed that CTPS-eGFP assemblies exhibited a range of movements—some were stationary, while others appeared to move more freely throughout the cell (Supplementary Movie 1). In other organisms, CTPS filaments assemble or disassemble in response to different environments. To understand if this is also true in mycobacteria, we exposed cells to a range of environmental conditions (Fig. 1d). To our surprise, in any condition that was associated with slowed or no growth, CTPS-eGFP was uniformly distributed throughout the cytoplasm. Transferring cells back to growth-permissive conditions resulted in rapid puncta formation (Fig. 1e). Together, these data reveal that, as in other species, mycobacterial CTPS forms filaments, and the presence of filaments is associated with growing *M. smegmatis*.

**Fig. 1:**
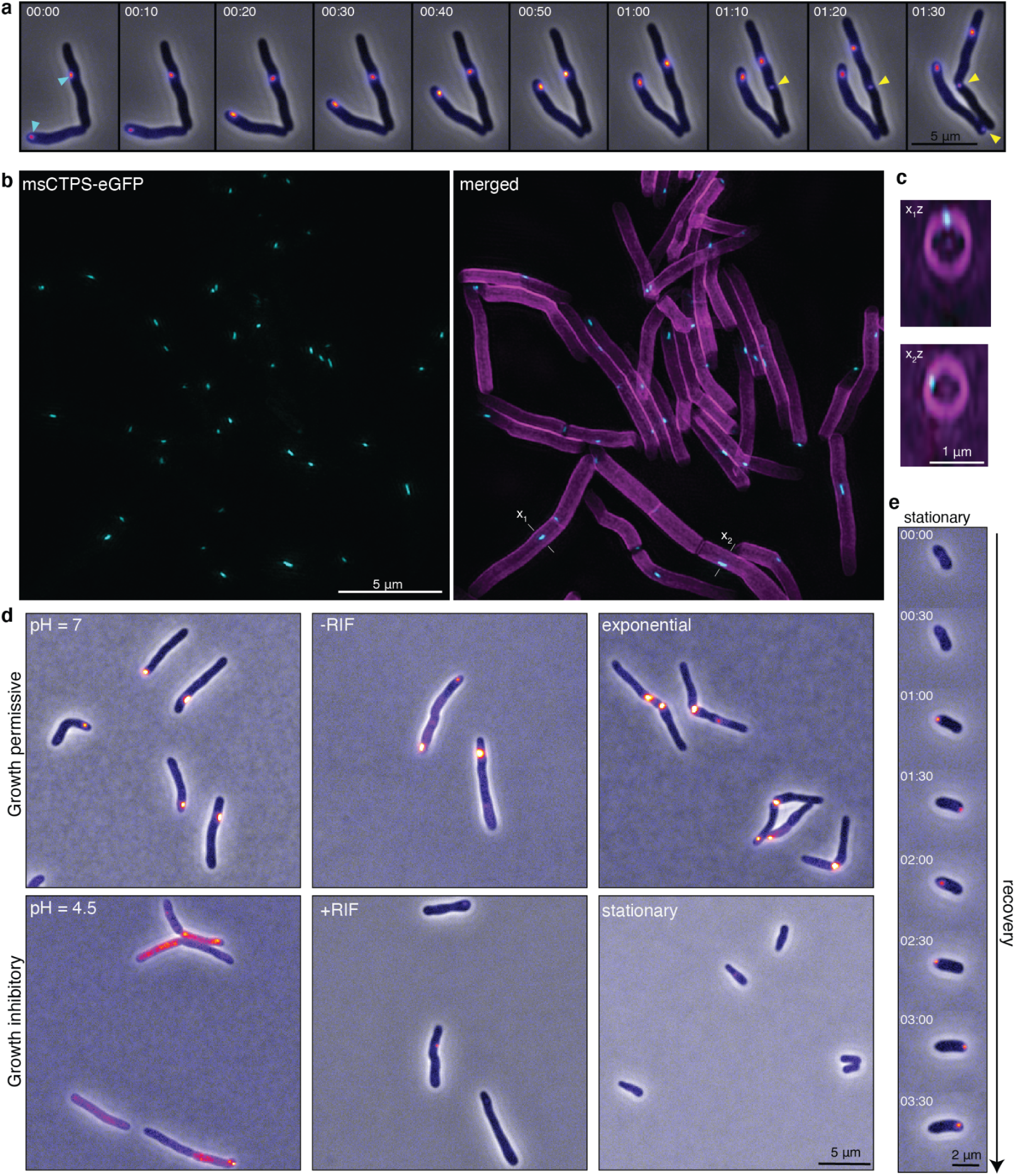
CTPS forms filaments in mycobacteria. **a,** Growing *M. smegmatis* cells expressing CTPS-eGFP from the endogenous locus are imaged over time by fluorescence and phase contrast microscopy. CTPS-eGFP is localized in discrete puncta (blue arrows). Puncta are mobile but often confined near the poles and septa. New puncta (yellow arrows) often appear at sites of division and new poles. **b,** Two-color three-dimensional structured-illumination microscopy (3D-SIM) of living CTPS-eGFP expressing *M. smegmatis* reveals that the puncta are small linear structures. Peptidoglycan is stained with a fluorescence d-amino acid (RADA) in the merged image. **c,** Two orthogonal views along the indicated axes (x_1_ and x_2_) show that CTPS-eGFP localizes near the membrane. **d,** CTPS-eGFP expressing cells were subjected to several environmental stresses that inhibit the growth of *M. smegmatis* (reduced pH, addition of rifampin, and growth to stationary phase). **e,** Stationary-phase cells were spotted on an agar pad containing fresh growth medium, and followed over time. Filaments form rapidly. Time is shown as hours:minutes in panels **(a)** and **(e)**.

### *M. tuberculosis* CTPS filaments have a distinct architecture

To further test the ability of mycobacterial CTPS to form filaments, we purified mtCTPS and investigated its ability to form filaments when isolated *in vitro*. Negative stain EM revealed that purified mtCTPS assembles into filaments when incubated with substrates UTP and ATP, but not with product CTP (Fig. 2a). Remarkably, from raw micrographs alone, it was immediately apparent that mtCTPS filaments did not resemble existing structures of CTPS filaments from other species, all of which have a characteristic appearance of stacked, X-shaped tetramers ^5,7–9,11^. In order to understand this differing architecture, we determined the cryo-EM structure of mtCTPS filaments in the presence of UTP and ATP at 3.2Å resolution (Fig. 2b and Fig. S2). The mtCTPS filament is composed of tetramers associated by residues upstream of the linker connecting the amidoligase and glutaminase domains (Fig. 2b,c). The amidoligase domains are positioned at the core of the filament, with glutaminase domains extending outwards (Fig. 2b). F280 from adjoining tetramers stacks at the center of the filament interface, with additional interactions formed between N277 and R274, as well as D282 and R38 (Fig. 2c). Immediately adjacent to this primary interface, there is a second, less well-resolved interface formed by the N-terminus, where residues 1-5 pack against the amidoligase domain of the neighboring tetramer (Fig. 2d). The active sites contained ADP and phosphorylated UTP, a reaction intermediate, suggesting that catalysis can occur within the filament (Fig. 2e). Similar intermediates were previously observed in a structure of active substrate-bound *Drosophila* CTPS ^5^.

**Fig. 2:**
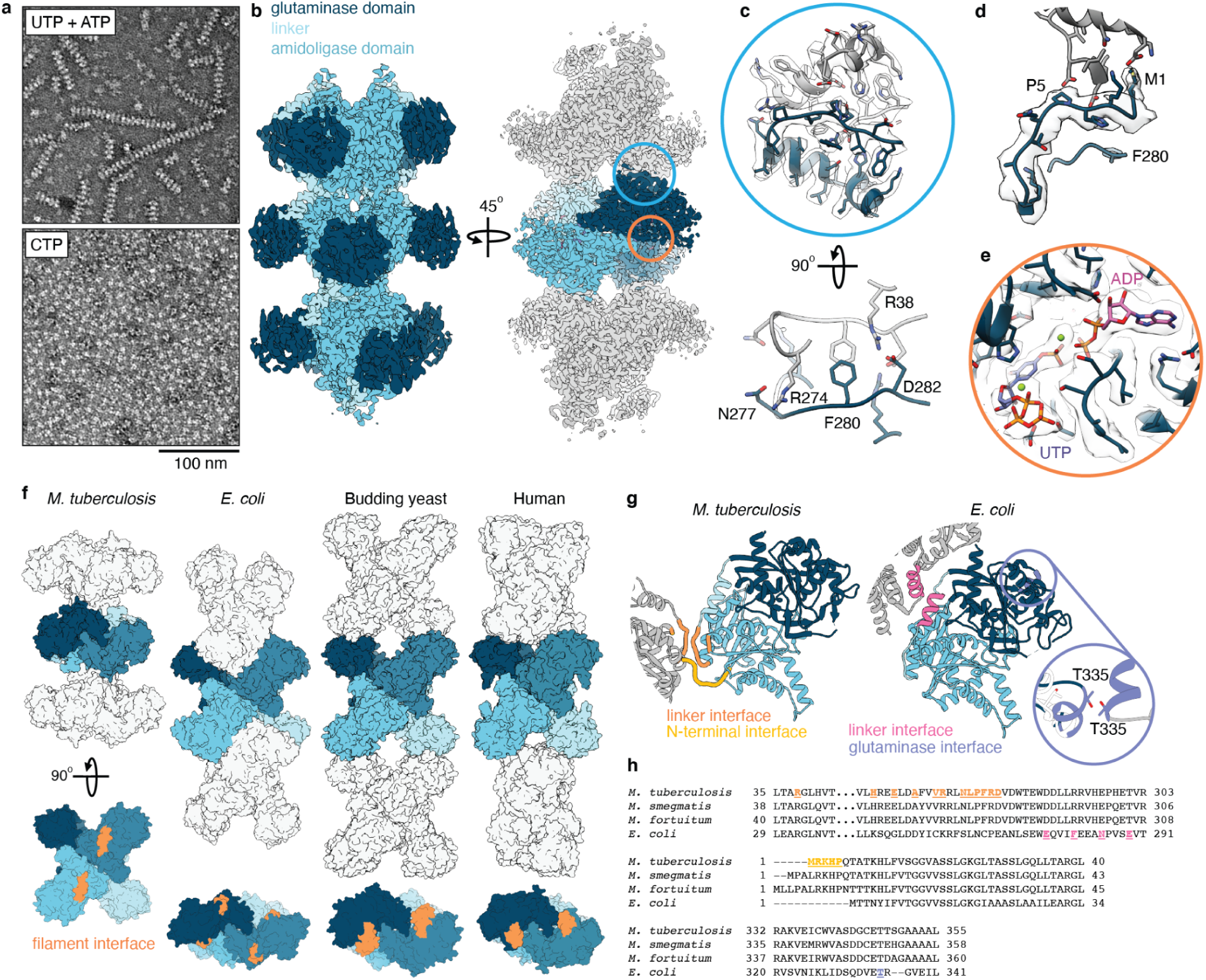
mtCTPS filaments have a distinct architecture. **a,** mtCTPS forms filaments when incubated with substrates UTP and ATP, but not with product CTP. **b,** Cryo-EM structure of the mtCTPS filament colored by domain (left), or with the central tetramer colored by monomer (right). **c,** Expanded view of the blue circle in **(b)**, showing the Cryo-EM map and atomic model at the filament interface. Lower panel shows the filament interface viewed down the helical axis, with interacting residues indicated. **d,** N-terminal filament contact site. **e,** Expanded view of the orange circle in **(b)**, showing phosphorylated UTP and ADP in the filament active site. **f,** Structures of CTPS filaments from various species. The central tetramer of each filament is colored by monomer, and filament interfaces are shown in orange. **g,** Comparison of the *E. coli* and *M. tuberculosis* CTPS filament interfaces. One monomer is colored by domain as in figure 1b, with adjoining monomers shown in gray. Filament interfaces are colored as indicated. The purple circle shows an expanded view of the *E. coli* glutaminase domain interface, overlaid with the *M. tuberculosis* structure in white. T335 at the *E. coli* interface is shown, together with the corresponding threonine residue in the *M. tuberculosis* structure. **h,** Sequence alignments comparing the filament interfaces in different bacteria, with important interface residues colored as in panel **(g)**.

The mtCTPS filament structure differs dramatically from other species’ CTPS filaments. Owing to the position of their filament assembly interfaces, mtCTPS tetramers pack in an orientation orthogonal to tetramers in human, *Drosophila*, yeast, or *E. coli* CTPS filaments, resulting in a distinct architecture (Fig. 2f). While both *E. coli* CTPS and mtCTPS filaments assemble via the region linking the amidoligase and glutaminase domains, the interfaces differ and are poorly conserved; the *E. coli* interface is on the helical portion of the linker, while the mtCTPS interface sits on a loop immediately upstream of this helix (Fig. 2g,h). *E. coli* CTPS forms an additional filament contact via the glutaminase domain. However, the fold differs in this region of the mtCTPS glutaminase domains which are, in any case, not positioned to interact with one another in the mtCTPS filament (Fig. 2b,g,h). The N-terminal mtCTPS filament interface is mediated by a short extension not present in other species (Fig. 2g,h and Fig. S3a). While both filament interfaces are highly-conserved amongst mycobacteria, the N-terminus is further extended in rapidly-growing species (Fig. S3a). We considered that this extension might sterically interfere with filament assembly. However, extending the mtCTPS N-terminus to resemble *M. fortuitum* CTPS (MLLPAL-mtCTPS), had no major effect on filament formation (Fig. S3b), overall suggesting that CTPS polymerization is likely conserved amongst mycobacteria. Further, deleting the five N-terminal residues of mtCTPS (ΔN-mtCTPS) reduced but did not abolish polymerization (Fig. S3b), indicating that the N-terminal contact plays a non-essential, stabilizing role in filament assembly.

### Clinical mutations affect mtCTPS polymerization and activity

Intriguingly, *pyrG* is under positive selection in clinical isolates of *M. tuberculosis*, suggesting that mutations in CTPS improve *M. tuberculosis* fitness in conditions relevant to TB drug sensitivity and/or pathogenesis ^25^. When we examined the selected mutations we noticed that several were at interesting positions on the protein structure. Specifically, we sought to understand the impact of mutations at the ATP binding site (P194S), and at the filament forming interface (H264R) (Fig. 3a). Thus, we purified mtCTPS-P194S and mtCTPS-H264R and compared their ability to form filaments with the wild-type enzyme. Strikingly, while mtCTPS-P194S retained its ability to form filaments, mtCTPS-H264R did not (Fig. 3b). To confirm this in the context of the cell, we mutated the homologous residues on the *M. smegmatis* chromosome using markerless single-stranded recombineering. In these strains, we again generated a chromosomal fusion to GFP, thereby fusing the mutant CTPS enzymes to GFP at the native locus. As predicted, msCTPS-P197S-GFP formed punctate assemblies while msCTPS-H267R-GFP exhibited uniform localization throughout the cell, confirming that the higher-order assemblies observed in cells indeed correspond to the filaments observed with purified protein (Fig. 3c).

**Fig. 3:**
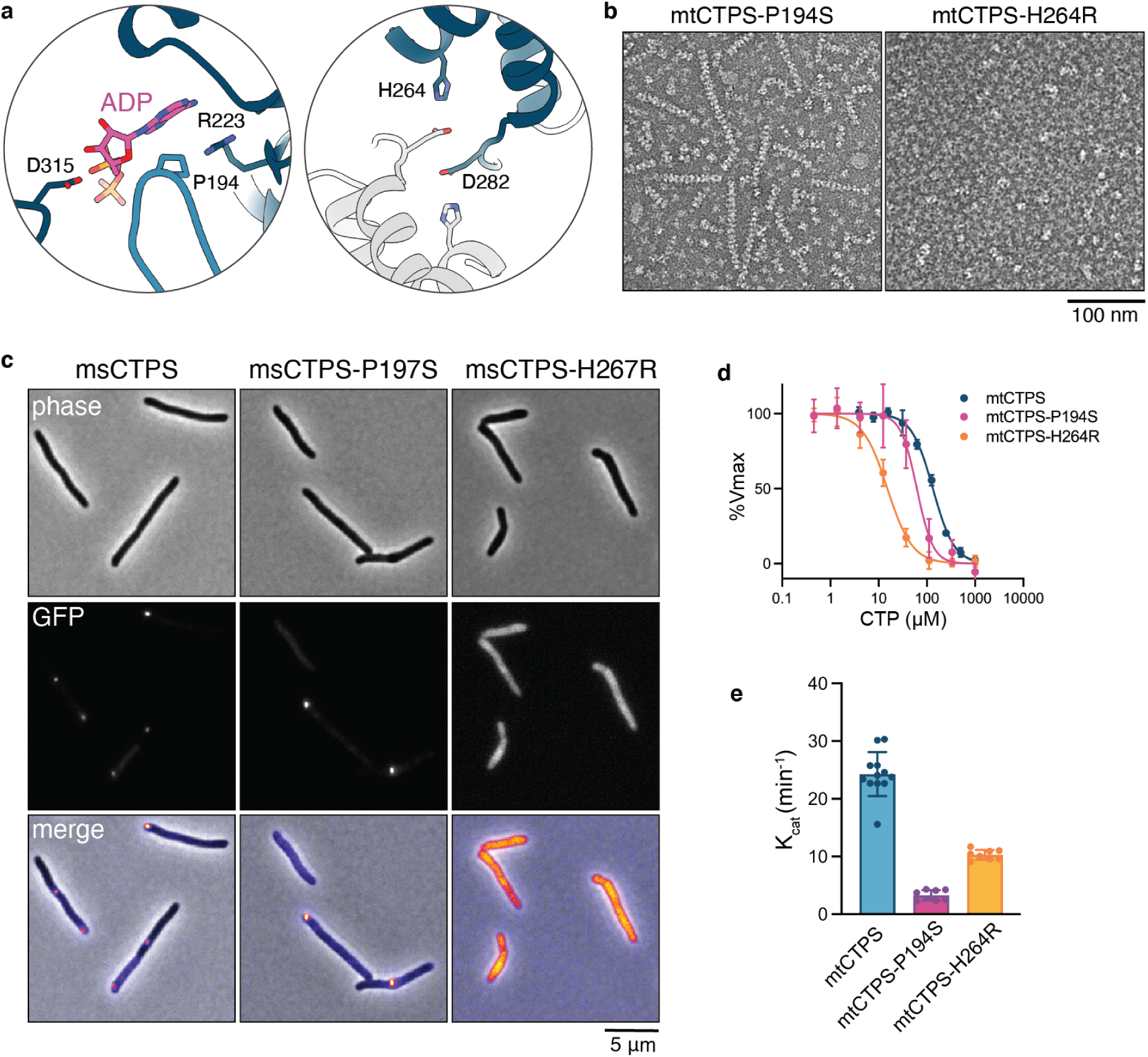
Clinical variants of mtCTPS exhibit reduced activity and polymerization. **a,** Positions of residues mutated in mtCTPS clinical variants, at the ATP-binding pocket (P194) and the filament forming interface (H264). **b,** Negative stain EM of mtCTPS clinical variants in the presence of UTP, ATP, and GTP. mtCTPS-H264R does not form filaments. **c,** Fluorescence imaging of GFP-tagged clinical variants of msCTPS in *M. smegmatis* cells. **d,** CTP inhibition curves for wild-type and mutant mtCTPS. Non-polymerizing mtCTPS-H264R has increased sensitivity to CTP inhibition. **e,** k_cat_ values for wild-type and mutant mtCTPS.

Next, we sought to understand the impact of these mutations on enzyme kinetics. mtCTPS-P194S exhibited values comparable to the wild-type enzyme for UTP, ATP, and glutamine substrate kinetics, GTP activation, and CTP inhibition (Fig. 3d and Fig. S4). By contrast, mtCTPS-P194S was overall less active, having a k_cat_ roughly 10-fold lower than wild-type mtCTPS (Fig. 3e). mtCTPS-H264R similarly had little effect on substrate or GTP affinities, and showed a more modest 2-fold reduction in k_cat_ (Fig. 3e and Fig. S4). However, mtCTPS-H264R was significantly more sensitive to CTP inhibition, having an IC_50_ for CTP nearly 10-fold lower than that of wild-type mtCTPS (Fig. 3d). Importantly, this reveals that mtCTPS filaments function to resist product feedback inhibition by CTP. Together, these results suggest that *M. tuberculosis* comes under selective pressure to reduce CTPS activity during infection, and achieves this via at least two distinct mechanisms: either by directly reducing the rate of catalysis or by preventing filament formation to increase sensitivity to CTP feedback inhibition. The selection against mtCTPS filament formation could also reflect a yet-unknown function for mtCTPS filaments that reduces fitness in some environmental niche encountered during infection or upon antibiotic challenge.

To better understand why mtCTPS filaments resist inhibition by CTP, we determined the cryo-EM structure of the CTP-bound mtCTPS tetramer at 3.6Å resolution (Fig. S5 and Fig. S6). CTP bound both the UTP and ATP binding sites of mtCTPS (Fig. S6a-c). Previous studies revealed that human and *Drosophila* CTPS similarly bind CTP at both these sites^4,5,7^, although with a different pose for the CTP base at the ATP site (Fig. S6d). CTP-bound mtCTPS exhibited the same characteristic conformational changes observed in existing product-bound CTPS structures: the glutaminase domains are rotated away from the amidoligase domains, while the tetramer interface is compressed to allow for CTP binding (Fig. 4a,b and Fig. S6e). Filaments cannot accommodate this product-bound conformation; compression of the tetramer upon CTP binding alters the relative positions of the two filament forming interfaces such that both can no longer interact simultaneously (Fig. 4c). mtCTPS filaments therefore appear to resist CTP feedback inhibition by stabilizing an active tetramer conformation incompatible with CTP-binding. This stabilization of the active, substrate-bound conformation may also account for the increased activity observed for wild-type mtCTPS over the non-polymerizing mtCTPS-H264R mutant.

**Fig. 4:**
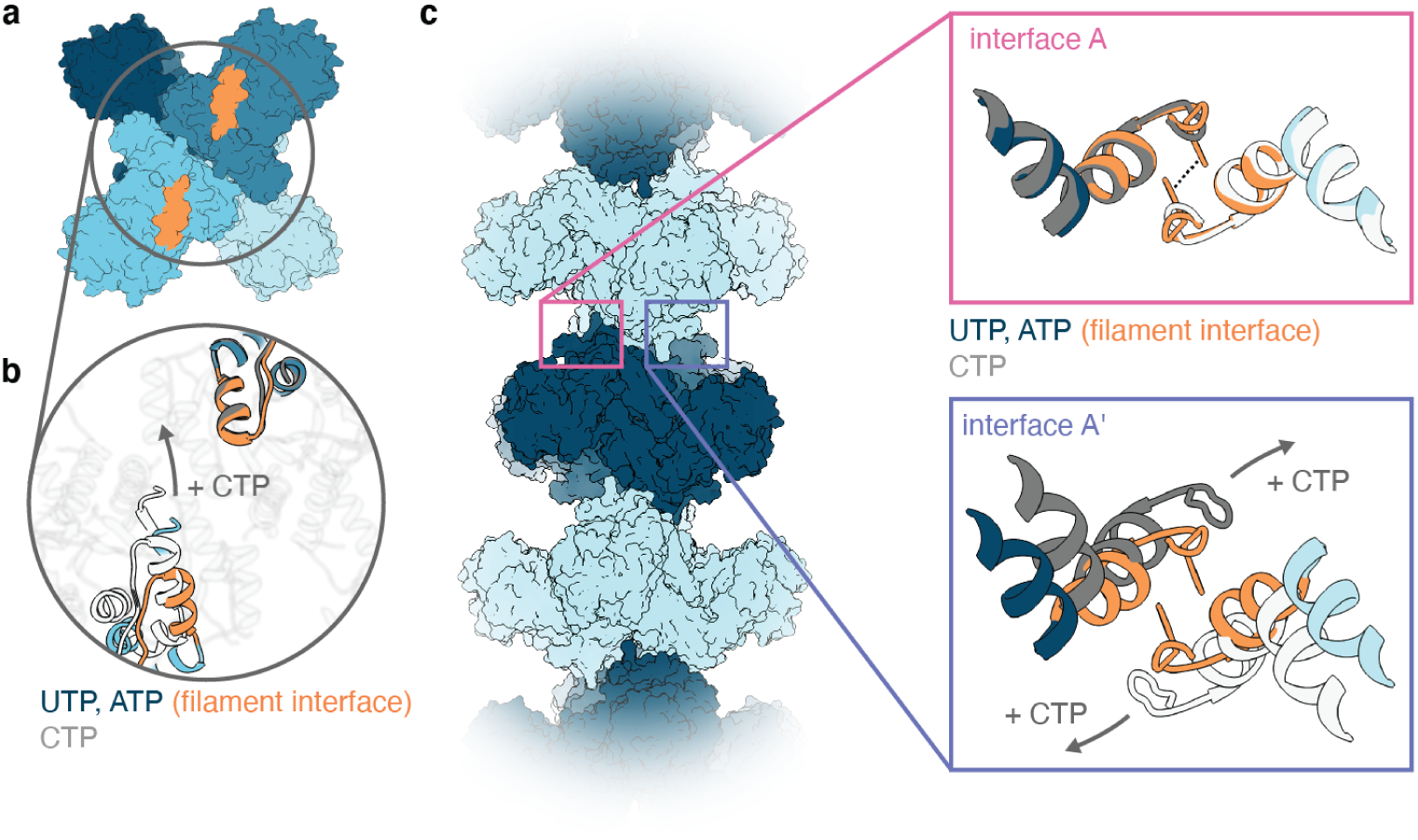
The CTP-bound conformation of mtCTPS disrupts filament formation. **a,** mtCTPS tetramer with positions of the filament interfaces shown in orange. **b,** Expanded view of the circle in **(a)**. CTP binding induces a compression of the tetramer interface, which alters the relative positions of the filament interfaces. **c,** mtCTPS filament with expanded views of the two symmetrical interfaces. Interaction between F280 is indicated by a dashed line. The substrate-bound mtCTPS filament structure and CTP-bound tetramer structure are aligned on filament interface A. CTP binding and resulting compression of the tetramer interface disrupts the interaction at filament interface A’.

### mtCTPS filaments flex to accommodate conformational changes

Although mtCTPS filaments stabilize the active enzyme conformation, a degree of flexibility was still apparent in the cryoEM structure, with the glutaminase domains poorly resolved in comparison to the amidoligase domains. Towards understanding this flexibility, we performed cryoSPARC 3D variability analysis (3DVA) with a low-pass filter resolution of 8Å, using a mask surrounding an individual tetramer. We then used Rosetta to relax atomic models into the maps representing the start and end of each 3DVA component (Fig. 5a,b). The movements observed by 3DVA represent the same general conformational changes expected when oscillating between the substrate- and product-bound states, though without reaching the fully compressed conformation observed when bound to CTP (Fig. 5c). 3DVA further revealed partial occupancy of the active sites; of the 4 active sites on each tetramer, only a subset were simultaneously bound to UTP. UTP binding induced the expected conformational changes in the enzyme: inward rotation of the glutaminase domain was coupled to extension of the tetramer interface, closing the active site around UTP (Fig. 5a,b). In the first 3DVA component, conformation and UTP occupancy was correlated within dimers, but anti-correlated across the tetramer interface, such that UTP bound only 2 active sites simultaneously (Fig. 5a,b). By contrast, in the second and third 3DVA components, conformation and UTP occupancy was correlated amongst 3 monomers, such that UTP was bound to either 1 or 3 active sites (Fig. S7). Overall, this suggests that mtCTPS may undergo partial-site reactivity, with substrate binding and catalysis occurring at only 1-3 active sites simultaneously. To understand how mtCTPS filaments can accommodate substantial flexibility without disassembling, we expanded 3DVA to include pairs of tetramers associated within filaments. This revealed that while tetramer interfaces extend and compress, complementary conformational changes in adjoining tetramers allows the filament interface to remain intact (Fig. 5d). These conformational changes may reflect motions required for the CTPS catalytic cycle, such that flexibility within filaments allows catalysis to occur without filament disassembly.

**Fig. 5:**
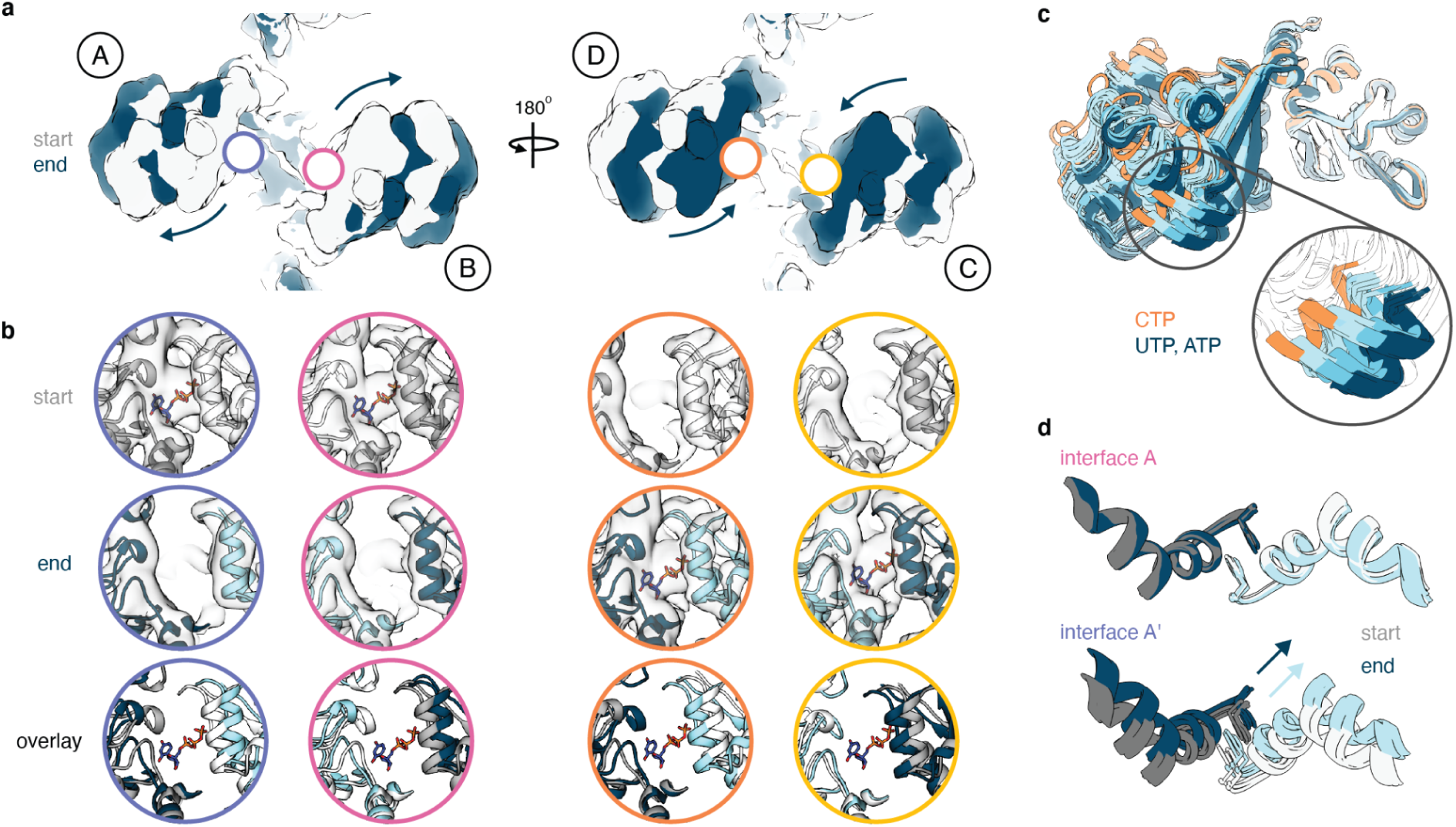
3D variability analysis of mtCTPS filaments. **a,** The first component from 3DVA performed on masked tetramers within mtCTPS filaments. Maps from the start and end of the 3DVA trajectory are shown in gray and blue, respectively. Monomers are labeled A-D. The glutaminase domains rotate back and forth, with outward rotation of the A and B monomers coupled to inward rotation of the C and D monomers. **b,** mtCTPS active sites corresponding to the colored circles in **(a)**. UTP binding is coupled with inward rotation of the glutaminase domains and closing of the active sites. **c,** Overlay of substrate-bound monomers from 3DVA (components 1-3) with product-bound mtCTPS. The substrate-bound monomers do not exhibit full rotation to the CTP-bound conformation. **d,** Results of 3DVA performed on pairs of tetramers within mtCTPS filaments, with the interface displayed as in Fig. 4c. Complementary conformational changes in adjoining tetramers allows filaments to remain intact despite flexibility at the filament interface.

### CTPS filaments are required for localization of the ParB partition complex

The ability of mtCTPS filaments to resist feedback inhibition by CTP could enable the highly localized production of CTP. The only protein known to be sensitive to CTP levels in mycobacteria is the CTPase ParB, which binds *parS* sites on the chromosome to mediate chromosome segregation. Upon binding CTP, ParB spreads along the DNA to form the partition complex that can be easily visualized as bright spots by fluorescent microscopy ^26,27^. To investigate if localized production of CTP affected the formation of the ParB partition complex in mycobacteria, we created *M. smegmatis* strains that express a second copy of ParB-mScarlet from the native promoter in wild type and CTPS mutant backgrounds. In wild type cells, consistent with its reported placement, ParB-mScarlet exists as 1 or 2 bright puncta near the ‘old pole’ at the beginning of the cell cycle (Fig. 6a,b). As the cell cycle progresses and the chromosome is duplicated and segregated, one ParB complex moves towards the other pole.

**Fig. 6:**
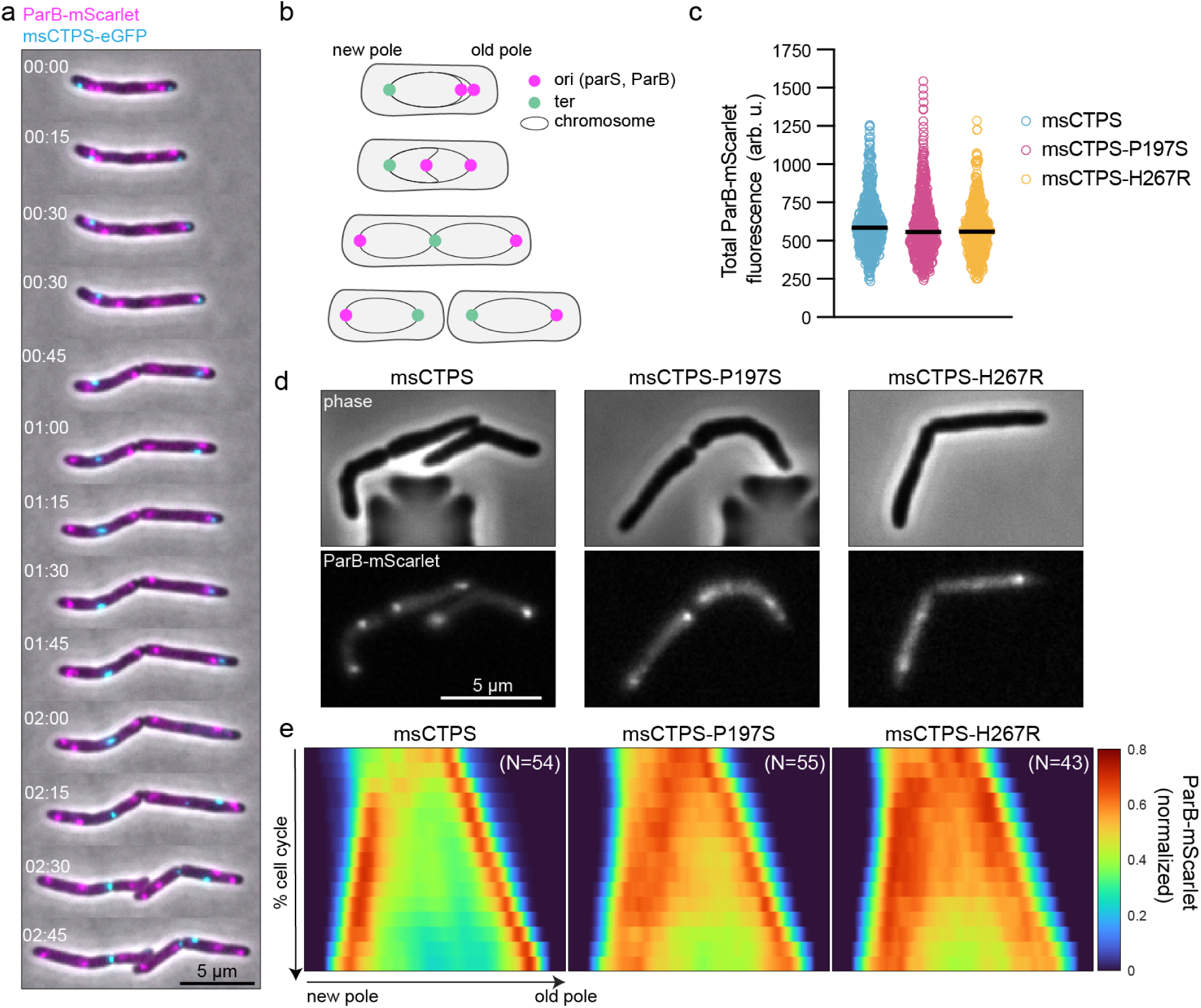
CTPS clinical variants exhibit abnormal ParB localization. **a,** *M. smegmatis* cells expressing CTPS-eGFP and ParB-mScarlet imaged over time. Time is shown as hours:minutes. **b,** ParB forms a CTP-dependent partition complex near the origin of replication and is needed for proper chromosome segregation. **c,** total amount of ParB-mScarlet in the indicated genetic backgrounds. **d,** ParB-mScarlet was visualized by phase and fluorescence microscopy in the indicated strains. **e,** To compare localization between strains individual cells were followed over time, from birth to division, and their fluorescence profiles were measured from new to old pole. Kymographs from the indicated number of cell cycles (N) were averaged together by 2D interpolation. The resulting ‘average kymograph’ represents the probability of finding ParB-mScarlet at a particular place in the cell at a particular time in the cell cycle.

Dual-color imaging of ParB-mScarlet and CTPS-eGFP revealed that the two complexes did not colocalize, but instead CTPS-eGFP often localized in the space between the ParB partition complex and the pole, which is presumably devoid of DNA (Fig. 6a). We next visualized ParB-mScarlet in cells expressing P197S or H267R variants of CTPS. In these mutant backgrounds, the total amount of ParB per cell is unchanged (Fig. 6c), but the overall intensity of the ParB partition complex is reduced, suggesting less efficient sliding of ParB along the DNA (Fig. 6d). Averaging ParB localization over several complete cell cycles revealed that, in cells expressing mutant CTPS the ParB complex is not as robustly formed at defined cellular localizations as in wild type cells, and its movement is less directed and more variable between cells (Fig. 6e). Together, these data suggest that one function of CTPS filaments is to provide localized CTP production for the robust formation of the ParB complex thereby promoting reliable chromosome segregation.

### Small-molecule inhibitors target the mtCTPS ATP-binding site

CTPS has been presented as a potential drug target in *M. tuberculosis*, primarily due to the essentiality of the *pyrG* gene, itself attributable to the requirement for CTP in the synthesis of various biomolecules. A number of small molecules from a GlaxoSmithKline antimycobacterial library were previously shown to target mtCTPS^23,28^. We selected two of these molecules, GSK1570606A and GSK735826A, and confirmed inhibition of purified mtCTPS. We obtained IC_50_ values similar to those previously reported, though with reduced maximum inhibition for GSK735826A (Fig. 7a). Contrary to the previous study, however, we observed selectivity for the *M. tuberculosis* enzyme; we did not observe inhibition of human CTPS1 or CTPS2 by either compound up to 100 µM concentration. Surprisingly, GSK735826A instead produced a 2-fold increase in CTPS1 activity with an EC_50_ value of 1.2 µM (Fig. 7a).

**Figure 7:**
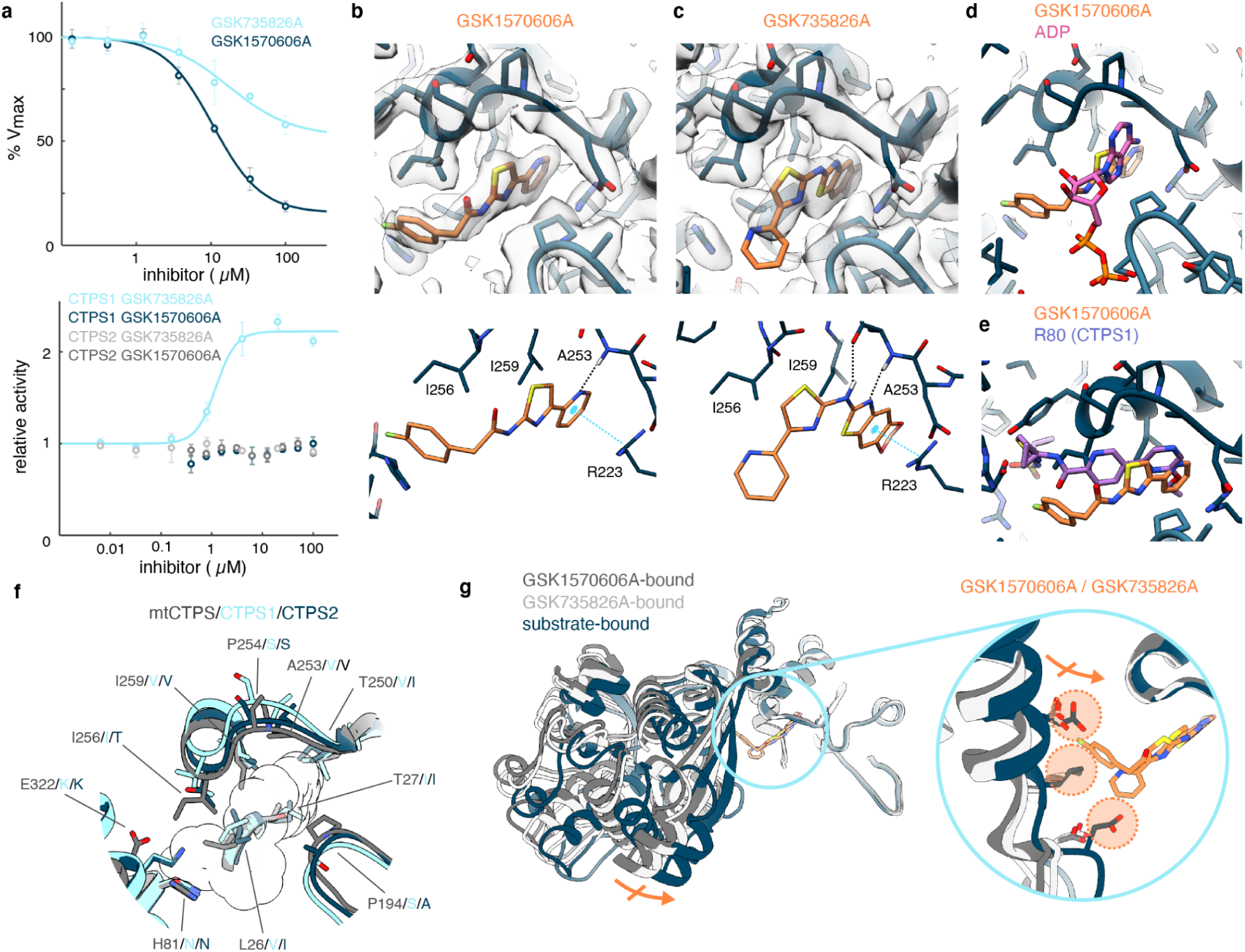
Inhibitors GSK1570606A and GSK735826A bind the ATP site of mtCTPS. **a,** Inhibition curves for GSK1570606A and GSK735826A against mtCTPS (top) or human CTPS1 and CTPS2 (bottom). **b,c,** Cryo-EM structures of mtCTPS bound to GSK1570606A **(b)** or GSK735826A **(c)**. Upper panels show cryo-EM maps and associated atomic models and lower panels show atomic models with important interacting residues. **d,** Overlay of GSK1570606A and ADP in the ATP-binding site of mtCTPS. **e,** Overlay of GSK1570606A and the human CTPS1 inhibitor R80. **f,** Comparison of the inhibitor binding pocket between mtCTPS, human CTPS1, and human CTPS2. Residues which differ are indicated. The volume occupied by GSK1570606A and GSK735826A is outlined. **g,** Comparison of the glutaminase domain rotation between the substrate-bound and inhibitor-bound mtCTPS structures. Structures are aligned on the amidoligase domain. The blue circle shows an expanded view of the inhibitor binding pocket, where rotation of the glutaminase domain to the substrate-bound conformation would produce clashes (orange circles) with GSK1570606A or GSK735826A.

To understand the mechanism of mtCTPS inhibition, we determined cryo-EM structures of mtCTPS bound to UTP, glutamine, and GSK1570606A or GSK735826A at 2.8Å resolution (Fig. 7b,c, Fig. S8 and Fig. S9). Consistent with the observation that these inhibitors are competitive with ATP binding^23^, they both bound sites overlapping with the ATP base, but with different poses (Fig. 7d). Notably, the terminal pyridine ring of GSK735826A was not well-resolved, suggesting a degree of flexibility in this region of the molecule (Fig. 7c).The orientation of the 4-(pyridin-2-yl)thiazole moiety common to both inhibitors is flipped between the structures, such that a different region of each inhibitor interacts with the backbone amide of A253 and forms pi-cation interactions with R223 (Fig. 7b,c). Similar interactions are observed between the pyrazine group of human CTPS inhibitors and the corresponding residues V247 and R217 in human CTPS1 and CTPS2. Although the human inhibitors thus function similarly by blocking ATP binding, the poses of the human CTPS and mtCTPS inhibitors differ substantially (Fig. 7e), while in both cases binding of UTP and Gln are not affected ^4^. A number of amino acid substitutions in the inhibitor binding pocket may account for the selectivity of GSK1570606A and GSK735826A for mtCTPS over the human enzymes (Fig. 7f). Further, while the human inhibitor-bound structures were in the same overall conformations as their respective substrate-bound structures^4^, the mtCTPS inhibitor-bound structures are in an intermediate conformation; the tetramer interfaces are in the extended conformation typical of the substrate-bound state, while complete inward rotation of the glutaminase domains is sterically prevented by inhibitor binding (Fig. 7g). Notably, *in silico* docking studies of GSK1570606A and GSK735826A by Esposito *et al.* ^23^ correctly predicted differing orientations of the 4-(pyridin-2-yl)thiazole moiety in the ATP site, though the docked poses differed overall from the cryo-EM structures. Together with the fact that CTPS is essential in mycobacteria, our demonstration that mtCTPS can be selectively targeted over the human enzymes, as well as the observation that mtCTPS may have a direct role in cell growth and division, further establishes CTPS as a bona fide drug target in *M. tuberculosis*.

## DISCUSSION

The mtCTPS filaments described here add to the already structurally diverse set of CTPS filaments from various species^5,7–9,11^. The apparently independent emergence of many distinct CTPS filament architectures, though initially surprising, is likely explained by the phenomenon described by the Levy laboratory: proteins have a propensity to form higher-order assemblies, with single surface mutations to hydrophobic residues often sufficient to produce polymerization^29,30^. This is particularly true for oligomeric proteins with dihedral symmetry, where filament-forming point mutations are presented on multiple symmetric surfaces, thus enhancing avidity—indeed, the majority of enzyme filaments are polymers of dihedral protein oligomers^31–34^. Consistently, although they differ greatly in their overall architectures, all CTPS filaments described to date are polymers of the conserved D2-symmetric CTPS tetramer. Further, all CTPS filament interfaces align along the dihedral symmetry axes of the tetramer, allowing for assembly into unbounded, linear polymers. The CTPS filament interfaces differ across species, but are generally distal to the more conserved active sites and allosteric regulatory sites, involving residues on the surfaces of the glutaminase and linker domains, as well as the N- and C-terminal tails^5,7–9,11^. These more variable regions are available for the evolution of filament forming interfaces that do not disrupt the core, conserved regulation of the CTPS tetramer, allowing polymerization-based regulation to layer on top of existing regulatory mechanisms.

Diversity in CTPS filaments architectures thus gives rise to diversity in regulation, with CTPS filaments from different species variously acting to increase or decrease activity, or to produce a more cooperative, switch-like response to CTP inhibition. mtCTPS filaments expand on this regulatory diversity by resisting feedback inhibition by CTP when compared with free tetramers. The substantial diversity in the structure and function of CTPS filaments suggests that CTPS polymerization arose independently multiple times, allowing for species-specific tuning of CTPS activity and regulation.

The cellular functions of enzyme filaments, though not completely defined, are varied. In many instances, functions appear to derive directly from the effects of polymerization on enzyme activity and consequent changes to metabolite levels^7,35–37^, though roles for enzyme filaments in cellular structure and organization have also been proposed^19^. While we do not yet know what functional role mtCTPS filaments play in mycobacterial physiology, our data provide some clues. The selection for mutants altered in filament formation in *M. tuberculosis* clinical isolates strongly suggests a functional role in mycobacterial physiology. Given our data, one likely role could be in the localized production of CTP needed for robust formation of the CTP-dependent ParB partition complex. Homologous mutations in *M. smegmatis* that are found to be positively selected for in *M. tuberculosis* alter enzyme kinetics and robust formation of the CTP-dependent partition complex needed for relatable and orderly segregation of the chromosome. Future research will be needed to understand the advantage that CTPS mutants confer to the pathogen, but defects in DNA metabolism have, in general, been linked to antibiotic tolerance and/or persistence ^38,39^.

Cell proliferation generally requires an increase in nucleotide availability, hinting at another possible function for mtCTPS filaments. For example, polymerization allows human IMP dehydrogenase 2 (IMPDH2) to resist feedback inhibition by its downstream product GTP, thus enabling expansion of cellular GTP pools^37^. Increased levels of GTP are required for cell proliferation and, consistently, IMDPH2 filaments have been observed in various proliferating cells^36,40–42^. Analogously, lymphocyte proliferation depends on increased levels of CTP. In humans, this is achieved by expression of the proliferative CTPS1 isoform, which resists feedback inhibition to enable expansion of CTP pools ^4,43^. The observations that mtCTPS filaments appear specifically during growth and resist feedback inhibition therefore suggests they may similarly function to expand CTP pools to support proliferation. The ability of *M. tuberculosis* to transition between proliferative and nonproliferative states is critical to its life cycle, warranting future studies into the relationship between mtCTPS polymerization and cell growth, as well as the selective advantages conferred by non-polymerizing mtCTPS mutations observed in clinical isolates.

## METHODS

### Bacterial strains and culture conditions

*M. smegmatis* mc^2^155 was grown in Middlebrook 7H9 broth supplemented with 0.05% Tween80, 0.2% glycerol, 5 gm/L albumin, 2 gm/L dextrose and 0.003 gm/L catalase or plated on LB agar. *Escherichia coli* DH5a cells were grown in LB broth or on LB agar plates. Concentrations of antibiotics used for *M. smegmatis* is as follows: 20 mg/ml zeocin, 25 mg/ml kanamycin, 50 mg/ml hygromycin, and 20 mg/ml nourseothricin. Concentrations of antibiotics used for *E. coli* used for *E. coli* is as follows: 40 mg/ml zeocin, 50 mg/ml kanamycin, 100 mg/ml hygromycin, and 40 mg/ml nourseothricin.

### Plasmid construction and strain generation

Strains used in this study are listed in Table 1.

#### Oligo-mediated Recombineering

For creating SNPs at the native *pyrG* locus, we used a strain containing a mutated hygromycin resistance cassette integrated at L5 as the base (pKM427). This strain was transformed with 1) a targeting oligo containing the mutation and 2) an oligo containing an SNP that repairs the mutated hygromycin cassette. Validation of these strains was performed by Hyg^R^ screening and by sequencing. The repaired HygR vector was swapped out using L5 allelic swapping and replaced with an empty vector marked with nourseothricin (NatR) resistance. To make eGFP fusions on these backgrounds or in wildtype *M. smegmatis* we used ORBIT ^44^ to remove the stop codon and insert eGFP in frame downstream of *pyrG*.

#### Integrating plasmids and associated strains

All plasmids were generated by isothermal assembly whereby insert and backbone shared 20-25 bp homology. These were then introduced into M. smegmatis by electroporation.

### Fluorescence Microscopy

#### Conventional

An inverted Nikon Ti-E microscope was used for the time-lapse and snapshot imaging. An environmental chamber (Okolabs) maintained the samples at 37°C. For snapshot imaging exponentially growing cells were immoblized under agar pads containing 1% UltraPure Agarose (ThermoFisher, #16500500) made with 7H9 base media lacking tyloxapol and ADC. For timelapse imaging, exponentially growing cells were cultivated in a B04 microfluidics plate from CellAsic (Millipore), continuously supplied with fresh 7H9 or minimal medium, where described, and imaged every 15 minutes using a 60x 1.4 N.A. Plan Apochromat phase contrast objective (Nikon). Fluorescence was excited using the Spectra X Light Engine (Lumencor), separated using single- or multi-wavelength dichroic mirrors, filtered through single bandpass emission filters, and detected with an sCMOS camera (ORCA Flash 4.0). Excitation and emission wavelengths are: GFP (Ex: 470; Em: 515/30), mScarlet (Ex: 555; Em: 595/40). To reduce phototoxicity exposure times were kept below 100ms for excitation with 470nm and below 300ms for mScarlet.

#### Three-dimensional structured illumination microscopy

Bacterial cells were immobilized under agar pads as described above. To visualize peptidoglcayn, 3uM of RADA was added an exponentially growing culture approximately 18 hours prior, and cells were washed in fresh 7H9 twice before imaging. 3D-SIM imaging shown in Figure 1 was performed on our custom live-cell 3D structured-illumination microscope based on previously published methods^45,46^. One major modification pertains to how the *s*-polarization of the illumination beams, essential for creating high-contrast SIM pattern and thus high-quality images, was maintained for all SIM angles. After a spatial-light modulator diffracts the incoming laser beam, the linearly polarized beams were first passed through a quarter-wave plate and became circularly polarized, and then were directed through a custom-made round-shaped compound linear polarizer (Laser Components) composed of 12 equally spaced sectors and a small central circular area. While the central area is non-polarizing, each sector linearly polarizes along the direction tangential to the midpoint of its arc^47^. Hence for each SIM orientation, the side beams are both linearly polarized perpendicular to the plane of incidence (i.e., *s*-polarized) and the central beam remained circularly polarized. Even though the central beam is not colinearly polarized with the side beams, we found that in practice this method is sufficient in creating high-contrast first-order 3D SIM illumination patterns. The main benefit is the zero time delay as opposed to the 10s of milliseconds delay typical for liquid crystal polarization rotating devices.

### Image Analysis

#### Segmentation

Phase contrast and fluorescence time-lapse images were analyzed in open-source image analysis software Fiji^48^. Images were processed with a custom pipeline using Fiji, iLastik^49^, and U-Net^50^. The U-Net plugin in FIJI was used as the platform for training and segmenting our phase-contrast microscopy images. Instead of using the raw images as the input, however, we first obtained the Hessian of Gaussian Eigenvalues (HoGE, with σ value of 0.7) of the raw images, either with the iLastik software or a custom MATLAB script translated from the relevant iLastik source code. Using the HoGE images as the input, we trained our U-Net model using about ten 2D images and their associated segmentation masks manually generated and the trained model was subsequently used in all our U-Net segmentation tasks. Output segmented files were then transferred to Fiji where we used the binary masks to manually annotate cells using the magic wand tool. Basic analysis was done with direct measurement of birth length, width, area, and mean fluorescent signal in Fiji.

#### Kymographs

Average kymographs shown in Figure 6e were made as previously described ^51,52^. In brief, using the segmented masks, cells at birth were manually selected. These were then input into a custom-matlab based program to follow cells over the course of the cell cycle, and extract fluorescence intensity averaged across the width of the cell. Average kymographs were generated by 2D interpolation over cell length and time.

### Purification of CTPS

The mtCTPS protein fused to either an N- or C-terminal 6His-TEV tag was expressed from the pET28b(+) vector in *E. coli* BL21(DE3) cells grown in LB media containing 50 µg/mL kanamycin. Expression was induced by addition of 1mM IPTG overnight at 25°C. Cells from 400 mL of culture were harvested by centrifugation at 5000 RPM at 4°C for 10 minutes, washed (50 mM Tris-HCl, 100 mM NaCl, pH 8.0), resuspended in lysis buffer (50 mM Tris-HCl, 200 mM NaCl, 10% glycerol, pH 8.0), then lysed using an Emulsiflex-05 homogenizer (Avestin, Ottawa, Canada) for 5 minutes at 15,000 PSI. Lysate was clarified by centrifugation at 14,000 RPM at 4°C using a Thermo Scientific Fiberlite F14-14 × 50cy rotor, then applied to a 5 mL HisTrap FF Crude column (GE) on an ÄKTA start chromatography system (GE) pre-equilibrated in column buffer (50 mM Tris-HCl, 500 mM NaCl, 10% glycerol, 25 mM imidazole, pH 8.0). The column was washed with 25 column volumes of column buffer, before eluting protein with 5 column volumes of elution buffer (50 mM Tris-HCl, 500 mM NaCl, 10% glycerol, 400 mM imidazole, pH 8.0). Fractions containing mtCTPS were pooled and dialyzed into storage buffer (20 mM Tris-HCl, 150 mM NaCl, 5% glycerol, 1 mM DTT, pH 8.0) using Snakeskin 3500 MWCO dialysis tubing (Thermo Scientific). Precipitated protein was removed by centrifuging at 14,000 RPM at 4°C using a microcentrifuge. Clarified mtCTPS was then flash-frozen in liquid nitrogen and stored at -80°C. The same procedure was used to purify the mtCTPS-D282R non-polymerizing mutant. Recombinant human CTPS1 and CTPS2 were purified as described previously ^4^.

### Negative Stain Electron Microscopy

To prepare samples for negative stain electron microscopy, protein was applied to glow-discharged, carbon-coated grids and stained with 0.7% uranyl formate. Samples were imaged using a Morgagni microscope (FEI) operating at 100 kV and 22,000× magnification on an Orius SC1000 CCD (charge-coupled device) camera (Gatan).

### Cryo-EM Sample Preparation And Data Collection

Cryo-EM samples were prepared by applying mtCTPS to glow-discharged CFLAT 2/2 holey-carbon grids (Protochips), blotting four times successively, then plunging into liquid ethane using a Vitrobot (ThermoFisher). Samples for the various cryo-EM structures included the following: substrate-bound filament, 19 µM mtCTPS (C-terminal His tag), 2 mM UTP, 2 mM ATP, 10 mM MgCl_2_; CTP-bound tetramer, 5 µM mtCTPS (N-terminal His tag), 2 mM CTP, 10 mM MgCl_2_; GSK1570606A-bound tetramer, 19 µM mtCTPS (N-terminal His tag), 2 mM UTP, 7.5 mM L-glutamine, 200 µM GSK1570606A, 10 mM MgCl_2_; GSK735826A-bound filament, 19 µM mtCTPS (C-terminal His tag), 2 mM UTP, 7.5 mM L-glutamine, 50 µM GSK735826A, 10 mM MgCl_2_. All samples were in buffer containing 20 mM Tris-HCl, 150 mM NaCl, 5% glycerol, and 1 mM DTT, pH 8.0. With the exception of the GSK1570606A-bound tetramer, all samples also included 5 mM octyl glucoside. Data for the substrate-bound filament and CTP-bound tetramer were collected on a Glacios (Thermofisher) equipped with a K2 summit direct detection camera (Gatan Inc.). Movies were acquired in counted mode with a pixel size of 1.16 Å/pixel with 50 frames and a total dose of 65 electrons/Å^2^. Data for the GSK1570606A- and GSK735826A-bound structures were collected on a Titan Krios (ThermoFisher) equipped with a K3 direct detection camera (Gatan Inc.), as well as a Quantum Gatan Imaging Filter energy filter (Gatan Inc.) operating in zero-loss mode with a 20-eV slit width. Movies were acquired in superresolution mode with a pixel size of 0.4215 Å/pixel with 75 frames and a total dose of 63 electrons/Å^2^. Leginon ^53^ software was used for automated data collection.

### Cryo-EM Data Processing

Cryo-EM data processing workflows are summarized in supplementary figures 2A, 4A, 6A, and 7A. Movies were aligned and dose-weighted using the Relion ^54^ implementation of MotionCor2 ^55^ or cryoSPARC ^56^ patch motion correction. Patch CTF estimation, particle picking, and 2D classification were performed in cryoSPARC. For the GSK1570606A sample, 2D classes representing short filaments did not contribute productively to structure determination, perhaps due to presence at the air-water interface, and were therefore excluded at the 2D classification stage; we subsequently found that addition of 5 mM octyl glucoside improved the distribution of filaments in ice, and therefore included octyl glucoside in other samples. 3D classification was performed using Relion. 3D refinement was performed using Relion auto-refine or cryoSPARC homogenous refinement, and in both cases beamtilt, anisotropic magnification, and defocus refinement were also performed. For datasets where MotionCor2 was used for movie alignment and dose-weighting, Bayesian polishing in Relion was also performed. 3D variability analysis of mtCTPS filaments was performed using cryoSPARC 3DVA ^57^, with a filter resolution of 8Å with results visualized using simple mode. Local resolution estimates were performed using Relion postprocessing.

### Model Building And Refinement

The crystal structure of *M. tuberculosis* CTP synthase (PDB 4ZDJ) was used as a starting model for refinement into cryo-EM maps, with ligands replaced as appropriate. Models were docked as rigid bodies into cryo-EM maps using Chimera ^58^ and refined using ISOLDE ^59^. Adaptive distance restraints were imposed within the glutaminase domains, restraining them to their initial geometries, owing to the lower resolution in this region of the maps. Ligands were also refined in ISOLDE and subsequently in Coot ^60^. Real space refinement was then performed in Phenix ^61^, with grid searches, Ramachandran restraints, and rotamer restraints disabled, and with starting model restraints enabled. Rosetta ^62,63^ was used to relax atomic models into maps from the first and last frame of each component from cryoSPARC 3D variability analysis.

### Adp-Glo Assays For Ctps Activity

ADP-Glo enzyme assays (Promega) were performed in buffer containing 50 mM K-HEPES pH 7.4, 10 mM MgCl2, 5 mM KCl, 5 mM octyl glucoside, and 1 mM DTT at room temperature in black, low-volume 384-well plates (Corning). CTPS reaction volumes were 6 µL. Substrate concentrations were 150 µM for UTP, ATP, GTP, and glutamine, except where varied to determine EC_50_ or K_m_ values. CTPS (500 nM) was incubated with UTP, ATP, and GTP for 4 minutes, after which glutamine was added to initiate the reaction. CTPS reactions were allowed to proceed for 4 minutes, terminated by addition of ADP-Glo reagent (6 µL), then incubated for 45 minutes prior to addition of kinase detection reagent (12 µL). After incubating for 45 minutes with detection reagent, luminescence was recorded on a Varioskan Lux (Thermo Scientific) microplate reader. For inhibition assays, CTP or small molecule inhibitors at various concentrations were added together with glutamine upon initiating the reaction.

### UV-Based Assays For CTPS Activity

UV-based assays for CTPS activity were performed with 500-2500 nM protein in the same buffer and using the same substrate concentrations described for the ADP-Glo assays. Reactions were performed in 96-well UV transparent plates (Corning) with a final volume of 90 µL at room temperature. CTPS was incubated with UTP, ATP, and GTP for 4 minutes, prior to addition of glutamine to initiate the reaction. Absorbance at 291 nm was measured over time using a Varioskan Lux (Thermo Scientific) microplate reader. CTP production was calculated using the change in extinction coefficient between UTP and CTP at 291 nm (1338 M^−1^cm^−1^)^64^ from the initial linear portion of the Abs_291nm_ versus time progress curves.

### Data Availability

Cryo-EM structures and atomic models have been deposited in the Electron Microscopy Data Bank (EMDB) and Protein Data Bank (PDB), respectively, with the following accession codes: EMD-42605, PDB: 8UV4 (mtCTPS filament bound to substrates); EMD-42611, PDB: 8UV8 (mtCTPS bound to CTP); EMD-42612, PDB: 8UV9 (mtCTPS bound to GSK1570606A); EMD-42613, PDB: 8UVA (mtCTPS bound to GSK735826A).

## Acknowledgments

We thank the Arnold and Mabel Beckman Cryo-EM Center at the University of Washington for the use of electron microscopes. This work was supported by NIH Grants R35 GM149542 and R01 AI153048 to J.M.K and Pew and Searle Scholar funds to E.H.R.

---

**Fig. S1.**
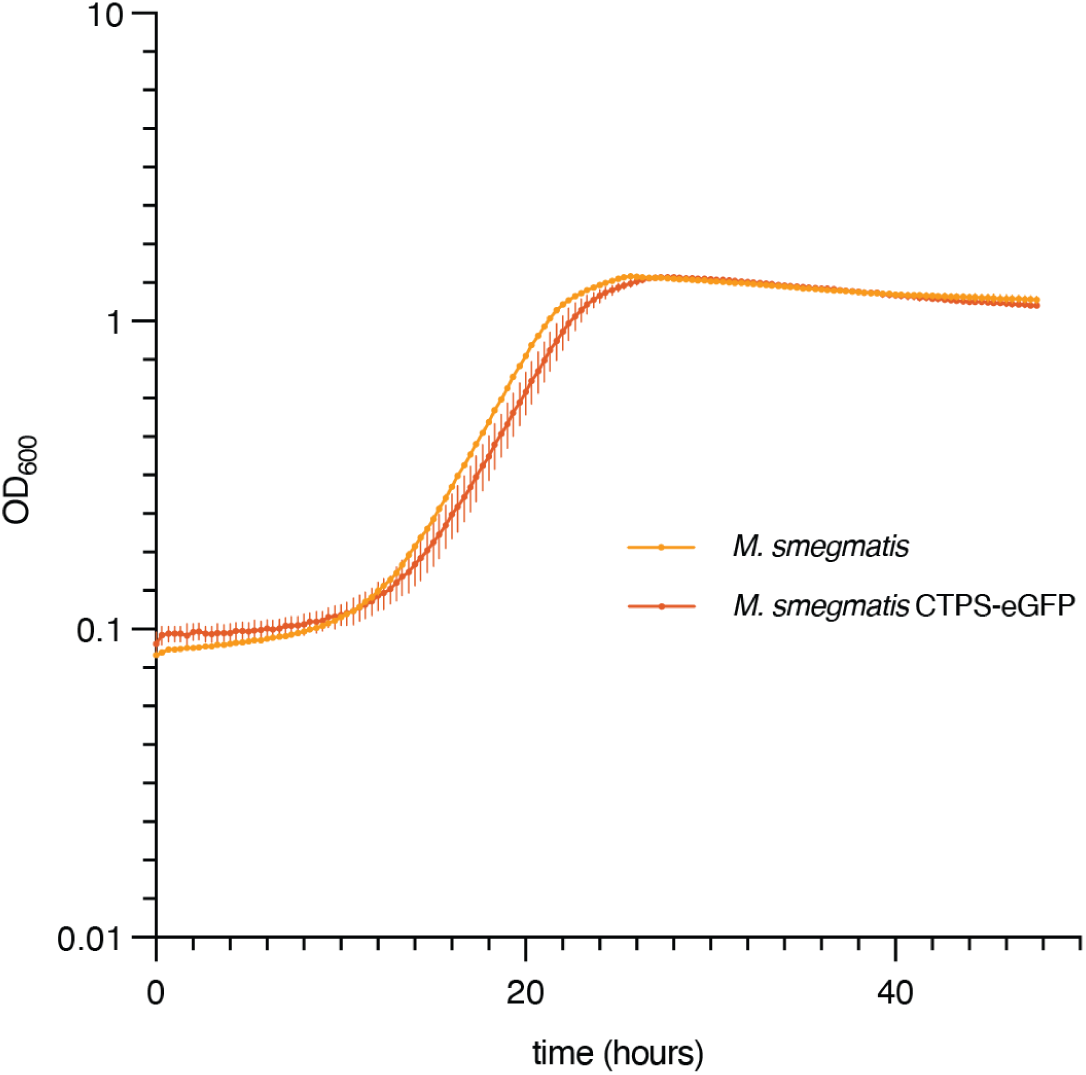
*M. smegmatis* expressing CTPS-eGFP exhibits normal growth. Growth of wild type *M. smegmatis* and *M. smegmatis* expressing CTPS-eGFP from its chromosomal locus was followed by optical density at 600 nm. The mean and standard deviation of three biological replicates is shown.

**Fig. S2.**
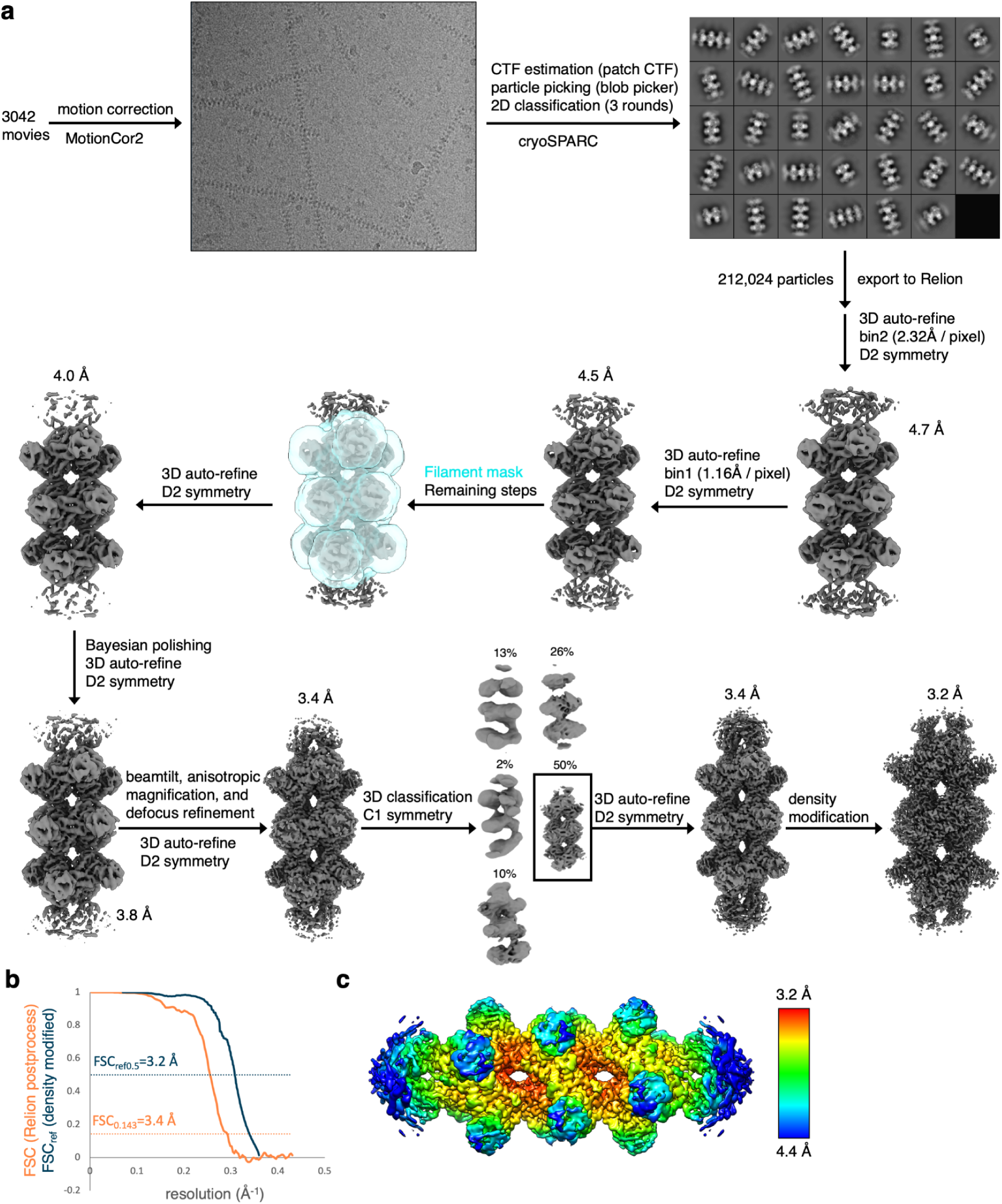
Cryo-EM data processing for mtCTPS filaments bound to ADP and phosphorylated UTP. **a,** Flowchart of cryo-EM data processing. **b,** Half-map FSC curve from Relion postprocessing (orange) and FSCref curve after density modification (blue) and corresponding resolution estimates. **c,** Relion local resolution estimate.

**Fig. S3.**
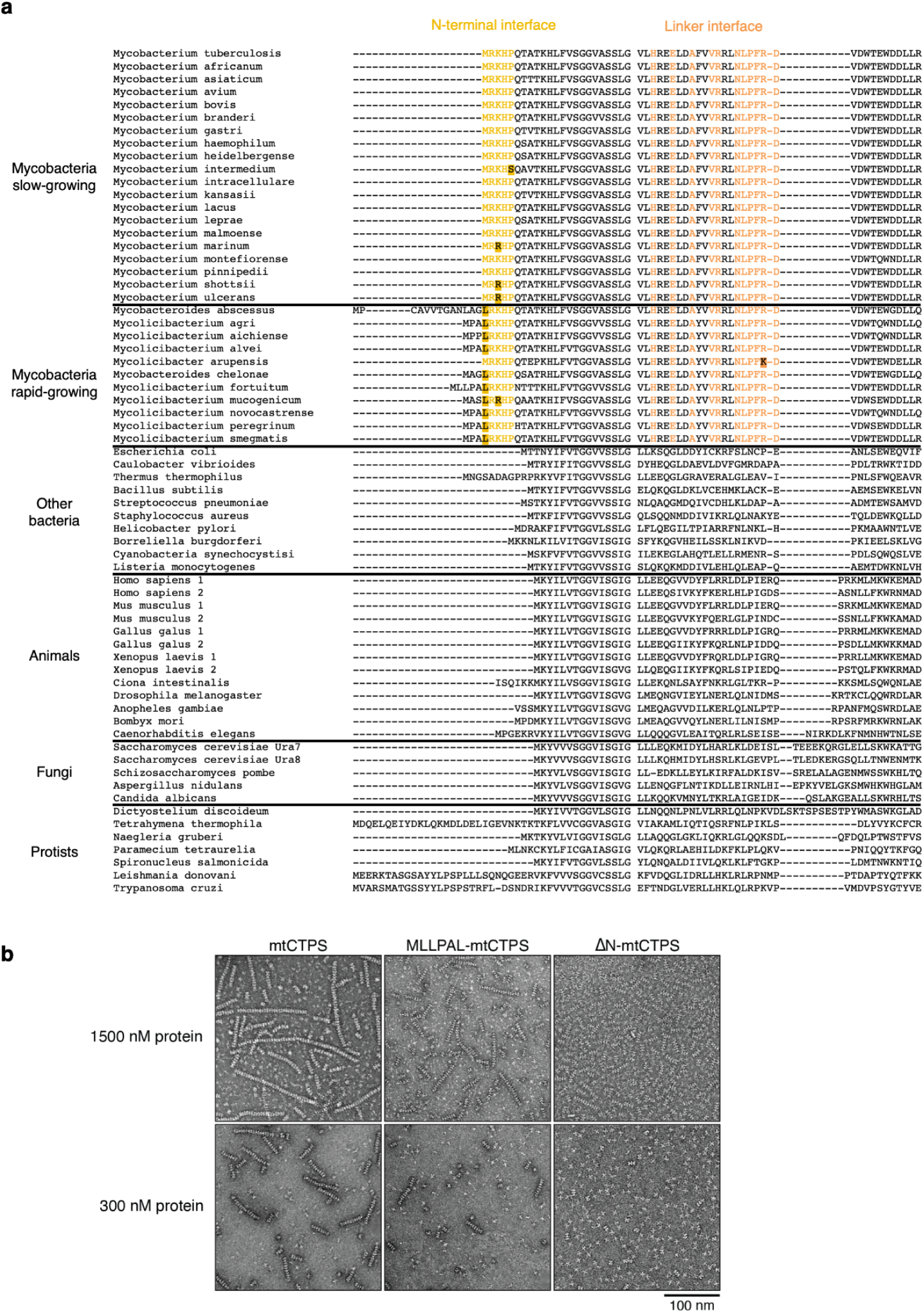
Mycobacterial CTPS filament-forming interfaces are not conserved in other species. **a,** Sequence alignments comparing the mtCTPS filament forming interfaces to other species. Conserved residues in the N-terminal and linker interface regions of various mycobacterial species are colored in yellow and orange, respectively. Residues in the interface regions differing from the *M. tuberculosis* sequence are highlighted. **b,** Negative stain EM images of wild-type and mutant mtCTPS at 300 nM or 1500 nM protein concentration in the presence of UTP, ATP, and GTP.

**Fig. S4.**
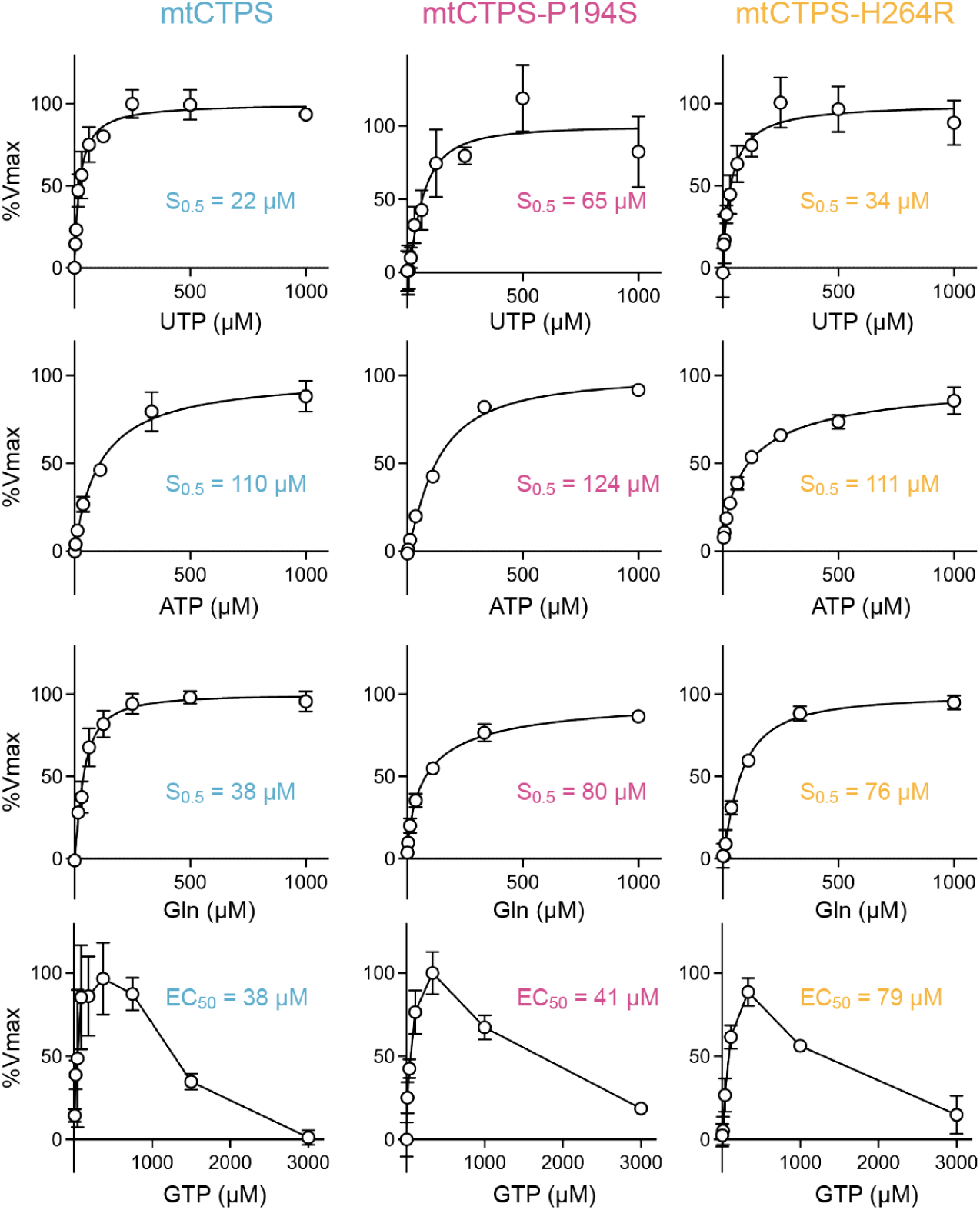
Enzyme kinetics for wild-type and mutant mtCTPS.

**Fig. S5.**
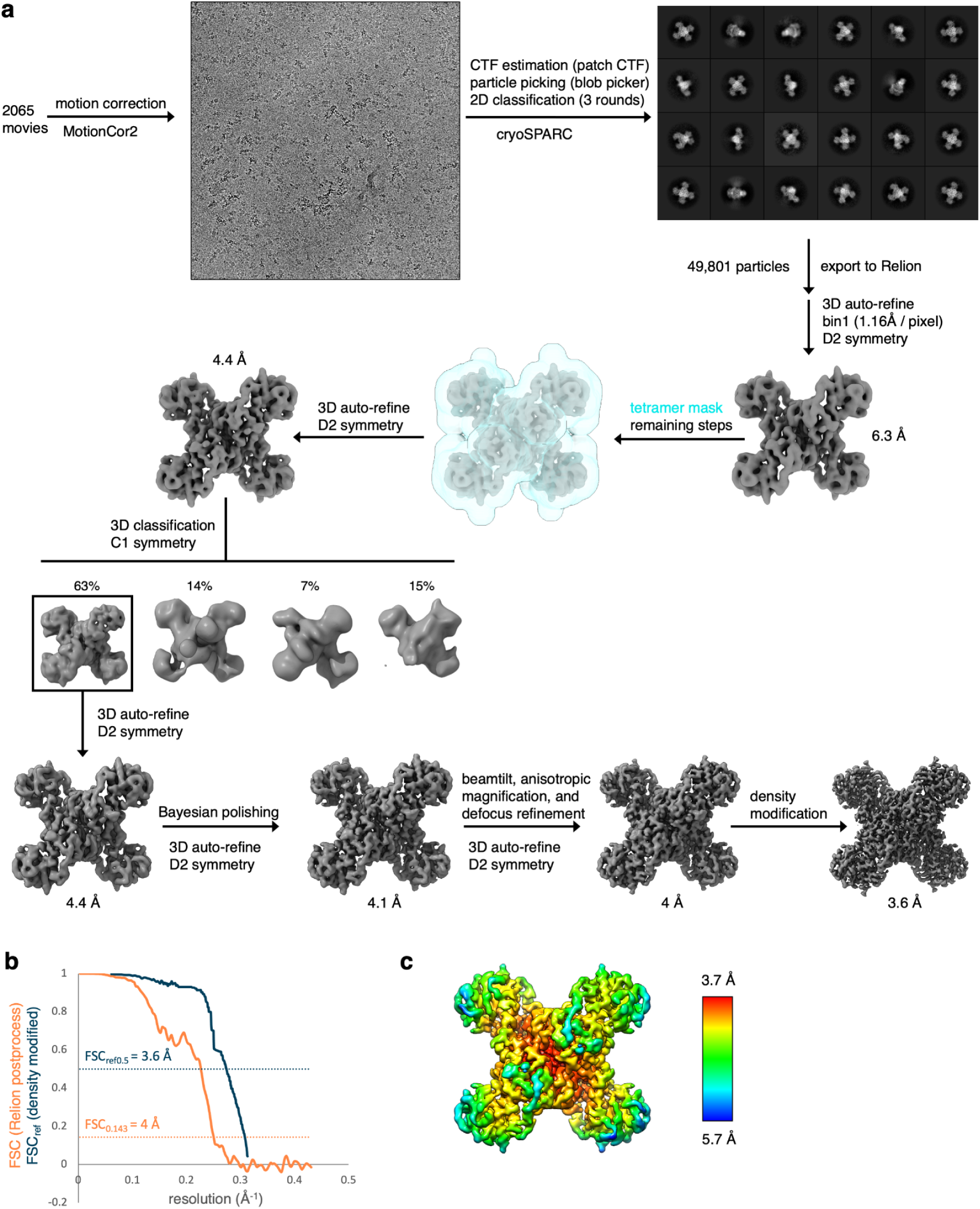
Cryo-EM data processing for mtCTPS tetramers bound to CTP. **a,** Flowchart of cryo-EM data processing. **b,** Half-map FSC curve from relion postprocessing (orange) and FSCref curve after density modification (blue) and corresponding resolution estimates. **c,** Relion local resolution estimate.

**Fig. S6.**
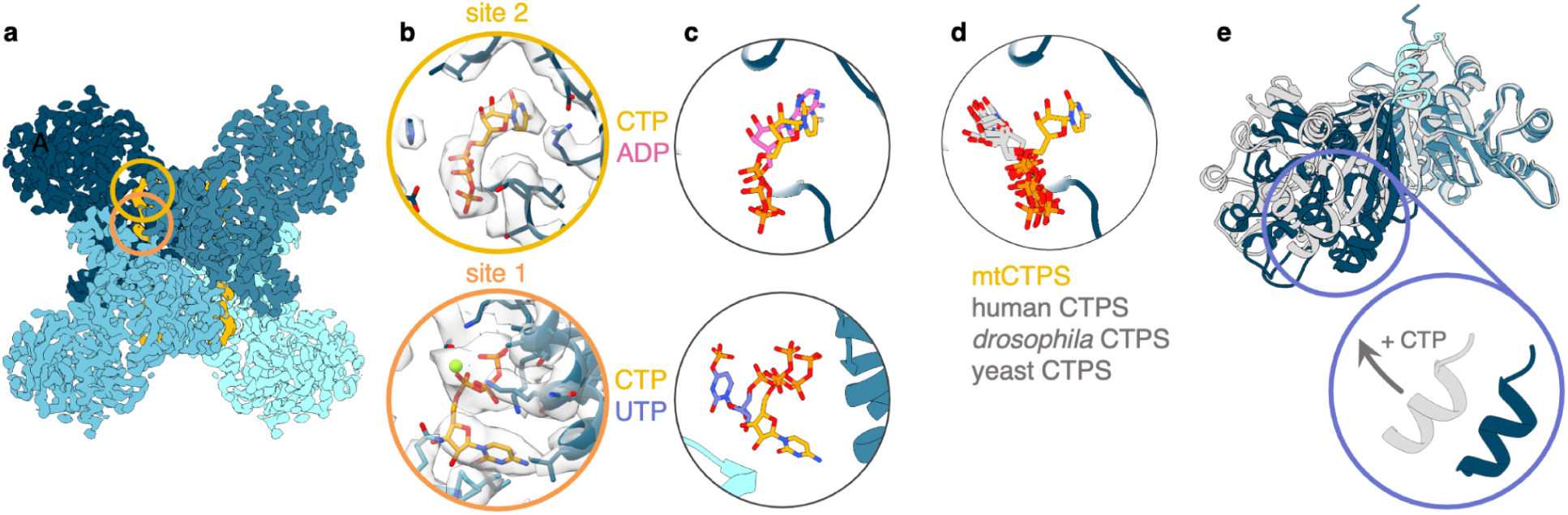
Structure of CTP-bound mtCTPS. **a,** Cryo-EM structure of CTP-bound mtCTPS at 3.6Å resolution. **b,** Expanded views of the circles in **(a)** showing CTP binding at sites 1 and 2. **c,** Overlays comparing CTP binding sites with ADP and phosphorylated UTP from the substrate-bound mtCTPS filament structure. **d,** Comparison of CTP binding at site 2 between mtCTPS and various eukaryotic CTPS structures. The orientation of the CTP base differs in mtCTPS. **e,** Overlay of substrate-bound (blue) and CTP-bound (gray) mtCTPS monomers, aligned on their amidoligase domains. The glutaminase domain is rotated away from the amidoligase domain in the CTP-bound structure.

**Fig. S7.**
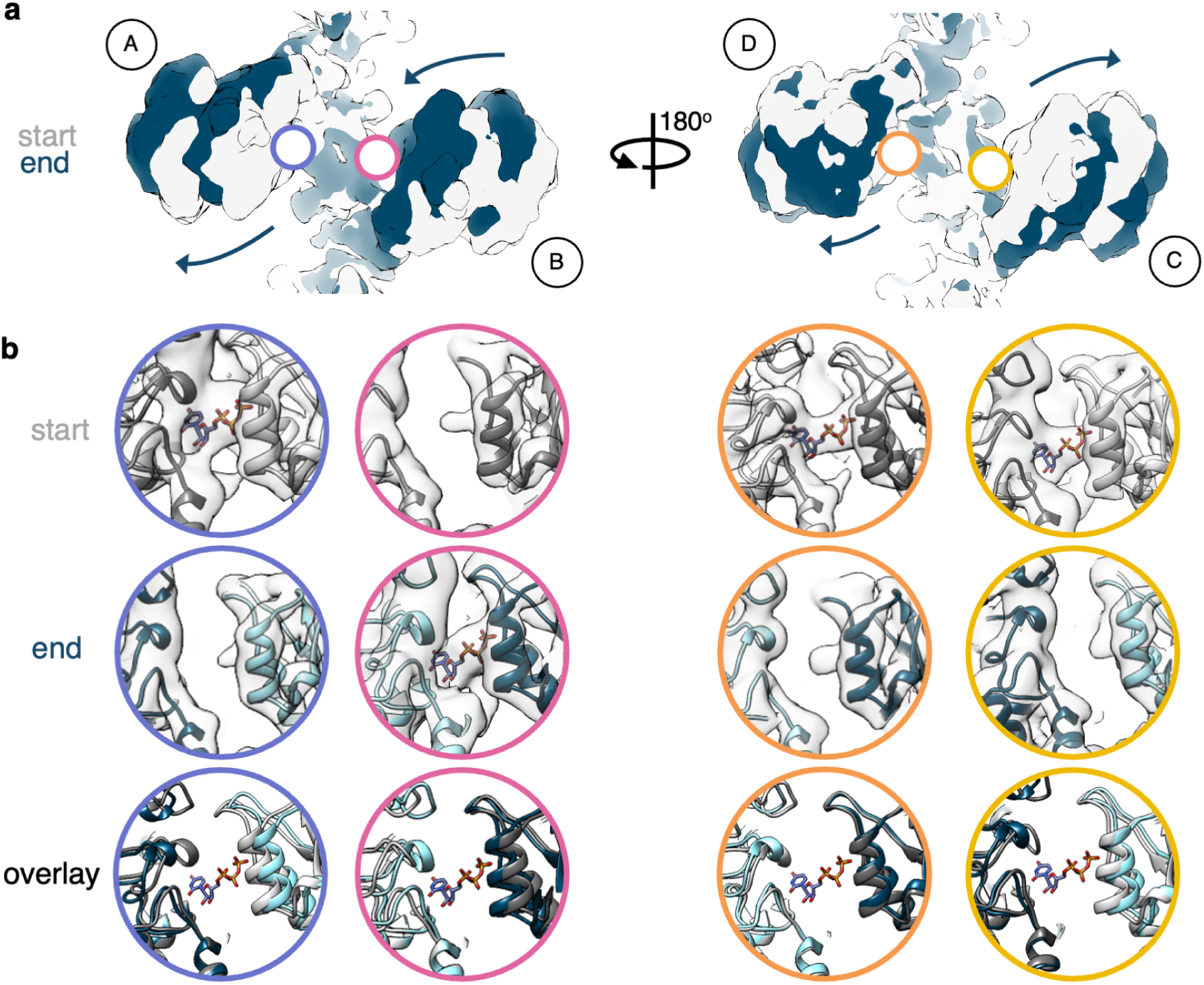
Second component from 3DVA performed on masked tetramers within mtCTPS filaments. **a,** Maps from the start and end of the 3DVA trajectory are shown in gray and blue, respectively. Monomers are labeled A-D. The glutaminase domains rotate back and forth. **b,** mtCTPS active sites corresponding to the colored circles in **(a)**. UTP binding is coupled with inward rotation of the glutaminase domains and closing of the active sites. Conformational changes in monomer D are less extreme compared with other monomers.

**Fig. S8.**
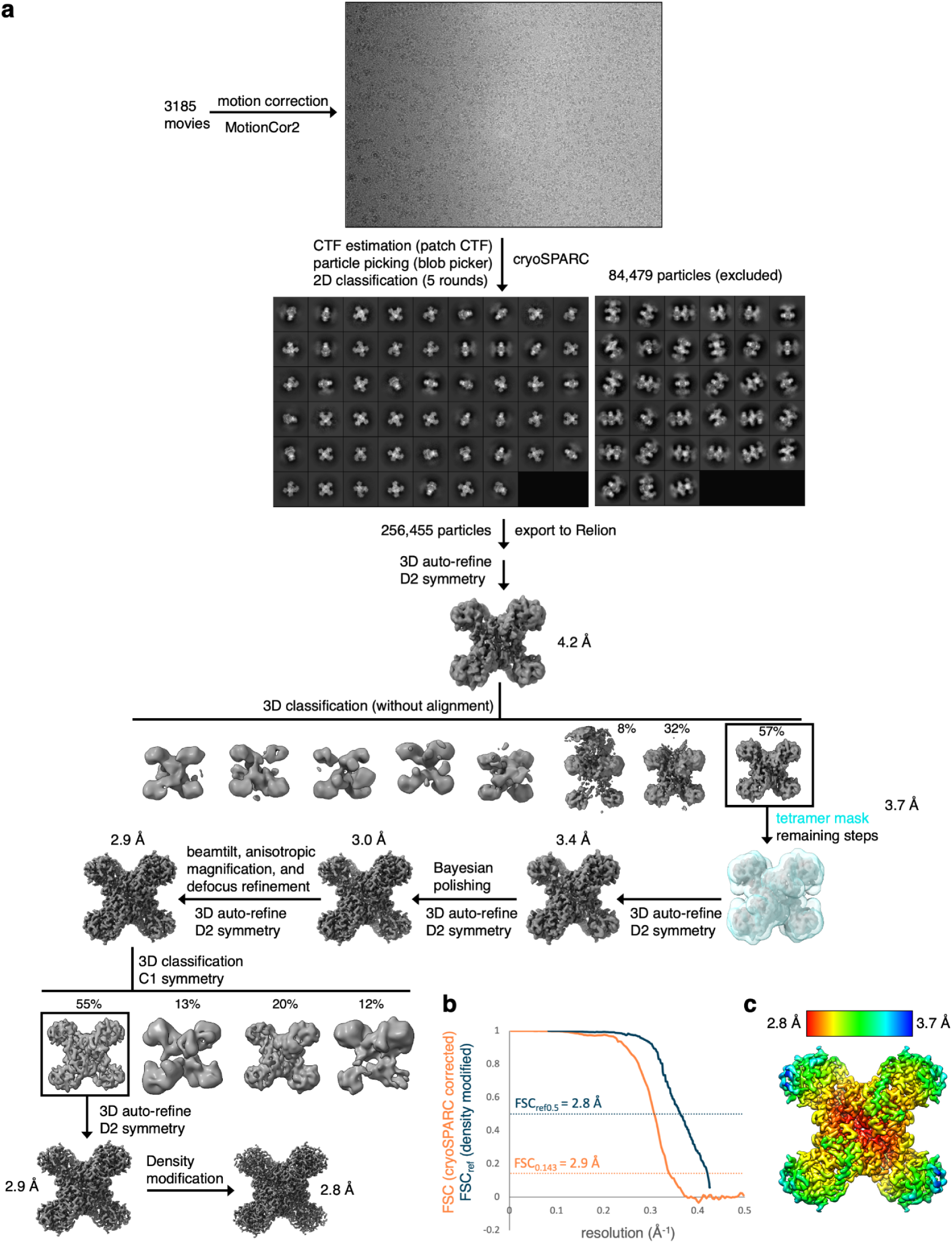
Cryo-EM data processing for mtCTPS bound to UTP, glutamine, and GSK1570606A. **a,** Flowchart of cryo-EM data processing. **b,** Half-map FSC curve from relion postprocessing (orange) and FSCref curve after density modification (blue) and corresponding resolution estimates. **c,** Relion local resolution estimate.

**Fig. S9.**
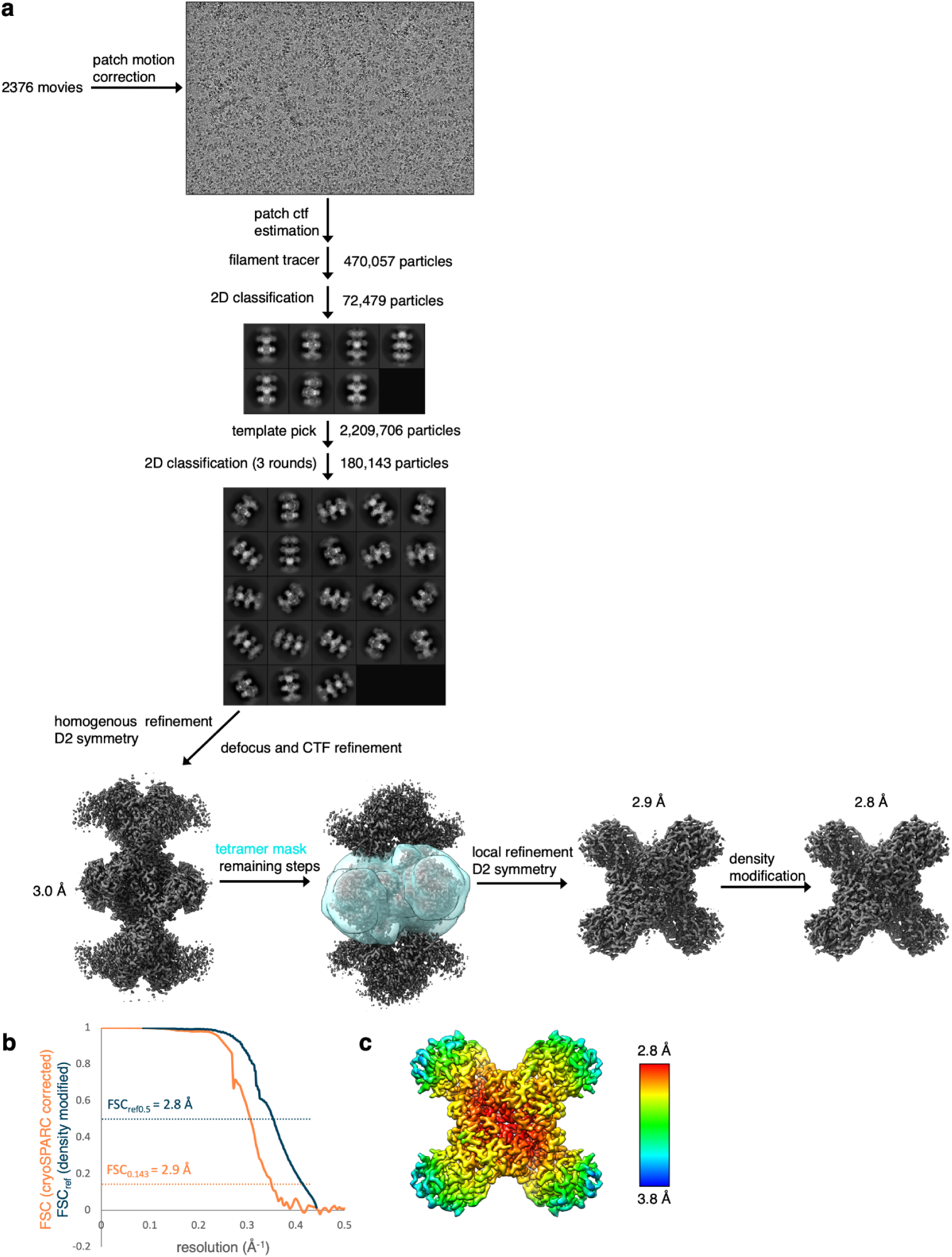
Cryo-EM data processing for mtCTPS bound to UTP, glutamine, and GSK735826A. **a,** Flowchart of cryo-EM data processing. All steps were carried out in cryoSPARC **b,** Half-map FSC curve (corrected) from cryoSPARC (orange) and FSCref curve after density modification (blue) and corresponding resolution estimates. **c,** Relion local resolution estimate.

**Table S1:** Strains used in this study.

| Reagent type (species) or resource | Designation | Source or reference | Identifiers | Additional Information |
| --- | --- | --- | --- | --- |
| strain (*Mycobacterium smegmatis*) |  |  | *Mycobacterium smegmatis* mc<sup>2</sup>155 | Wild type *M. smegmatis* |
| strain (*M. smegmatis*) | YL375 | This work | mc2155 *pyrG*::eGFP-pKM468 L5::NatR empty vector | Localization of CtpS WT. |
| strain (*M. smegmatis*) | YL440 | This work | mc2155 *pyrGP197S*::eGFP-pKM468 L5::NatR empty vector | Oligo recombineered SNP strain of the chromosomal *pyrG* with the P197S mutation and in-frame integration of eGFP |
| strain (*M. smegmatis*) | YL441 | This work | mc2155 *pyrGH267R*::eGFP-pKM468 L5::NatR empty vector | mc2155 *pyrGH267R*::eGFP-pKM468 L5::NatR empty vector |
| strain (*M. smegmatis*) | YL441 | This work | mc2155 L5::NatR empty vector | A genetically matched control strain for YL372 |
| strain (*M. smegmatis*) | YL418 | This work | mc2155 *pyrG*::eGFP-pKM468 L5::pNative-ParB-FLAG-mScarlet | Localization of CTPS and ParB |
| strain (*M. smegmatis*) | YL394 | This work | mc2155 L5::pNative-ParB-FLAG-mScarlet | Localization of ParB in WT |
| strain (*M. smegmatis*) | YL438 | This work | mc2155 *pyrG P197S* L5::pNative-ParB-FLAG-mScarlet | Localization of ParB in P197S mutant |
| strain (*M. smegmatis*) | YL439 | This work | mc2155 *pyrG H267R* L5::pNative-ParB-FLAG-mScarlet | Localization of ParB in H267R mutant |

**Table S2.**
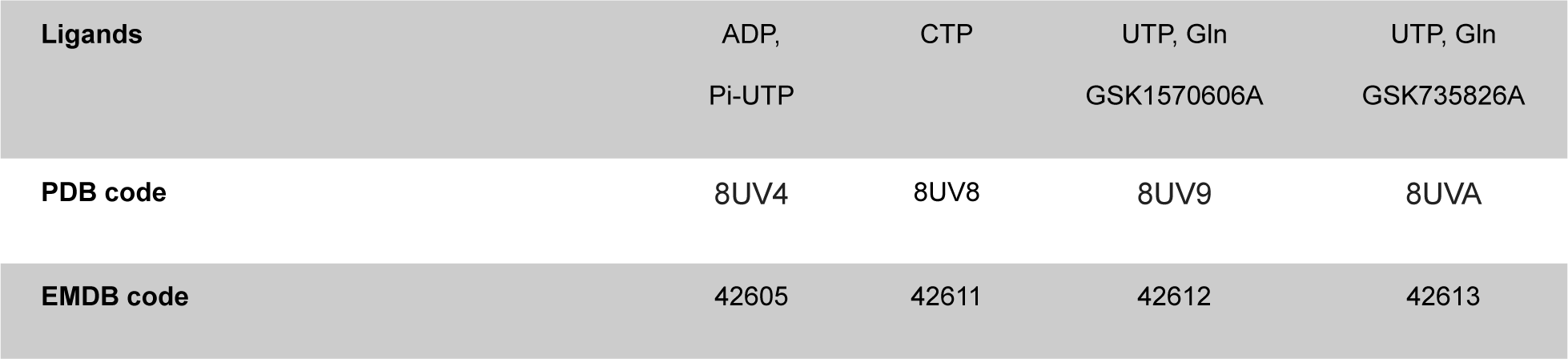
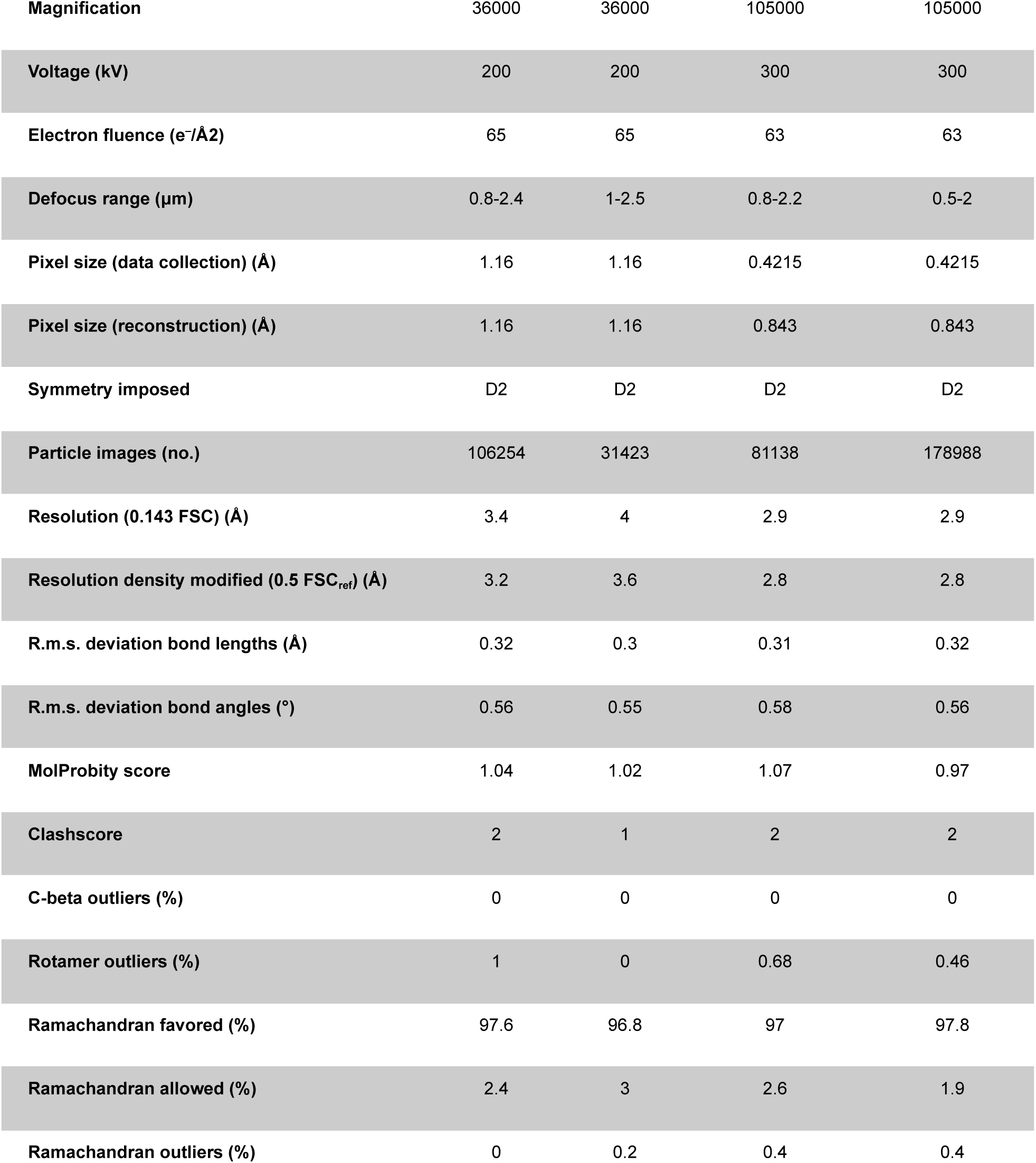
Cryo-EM data collection, refinement and validation statistics.

