## Supplementary figures and images for "Evolutionarily divergent *Mycobacterium tuberculosis* CTP synthase filaments are under selective pressure"

### Supplementary Movie 1

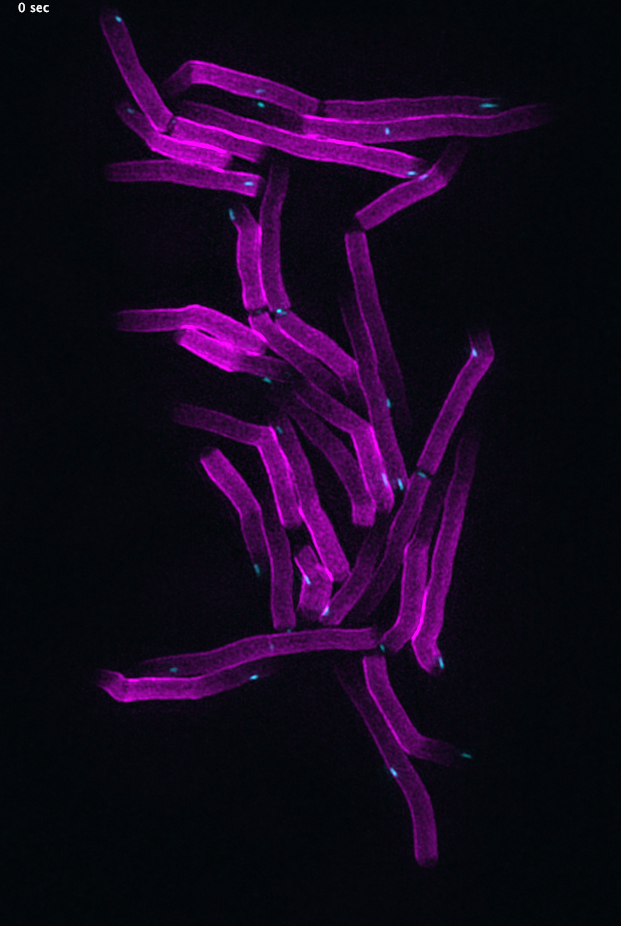
